# Neurodevelopmental deficits and cell-type-specific transcriptomic perturbations in a mouse model of *HNRNPU* haploinsufficiency

**DOI:** 10.1101/2020.05.01.072512

**Authors:** Sarah A. Dugger, Ryan S. Dhindsa, Gabriela De Almeida Sampaio, Andrew K. Ressler, Elizabeth E. Rafikian, Sabrina Petri, Verity A. Letts, JiaJie Teoh, Junqiang Ye, Sophie Colombo, Mu Yang, Michael J. Boland, Wayne N. Frankel, David B. Goldstein

## Abstract

Heterozygous *de novo* loss-of-function mutations in the gene expression regulator *HNRNPU* cause an early-onset developmental and epileptic encephalopathy. To gain insight into pathological mechanisms and lay the potential groundwork for developing targeted therapies, we characterized the neurophysiologic and cell-type-specific transcriptomic consequences of a mouse model of *HNRNPU* haploinsufficiency. Heterozygous mutants demonstrated neuroanatomical abnormalities, global developmental delay, impaired ultrasonic vocalizations and increased seizure susceptibility, thus modeling aspects of the human disease. Single-cell RNA-sequencing of hippocampal and neocortical cells revealed widespread, yet modest, dysregulation of gene expression across mutant neuronal subtypes. We observed an increased burden of differentially-expressed genes in mutant excitatory neurons of the subiculum—a region of the hippocampus implicated in temporal lobe epilepsy. Evaluation of transcriptomic signature reversal as a therapeutic strategy highlights the potential importance of generating cell-type-specific signatures. Overall, this work provides insight into *HNRNPU*-mediated disease mechanisms, and provides a framework for using single-cell RNA-sequencing to study transcriptional regulators implicated in disease.

## Introduction

Given their functional complexity, high metabolic demands and extensive diversity, it is unsurprising that neurons rely particularly on the strict regulation of gene expression. In fact, mutations in genes that cause gene expression dysregulation, including chromatin modifiers^1,2^, transcription factors^3,4^ and RNA-binding proteins^5,6^, are a well-described cause of neurodevelopmental disease, including epilepsy and autism. Considering that mutations in this class of molecules often lead to the dysregulated expression of thousands of genes within vulnerable cell types, pinpointing therapeutically tractable disease mechanisms is especially challenging. These pleiotropic effects necessitate the use of high-resolution phenotyping assays. One powerful approach is single cell RNA-sequencing (scRNAseq), which allows for the identification of cell-type-specific gene expression changes in disease-associated tissues. Here, using scRNAseq, we explore a novel, transcriptome-guided precision medicine approach on a genetic mouse model of *HNRNPU* haploinsufficiency.

*HNRNPU* (heterogeneous nuclear ribonuclear protein U) encodes a ubiquitously-expressed, DNA- and RNA-binding protein that localizes to the nucleus^7,8^, where it mediates gene expression through transcription initiation and elongation^9–14^, pre-mRNA processing^15,16^ and chromatin organization^17,18^. We and others have reported *de novo* loss-of-function variants^19–23^ and microdeletions^24,25^ encompassing *HNRNPU* in pediatric patients with a severe, and often treatment refractory, developmental and epileptic encephalopathy (DEE) characterized by early-onset epilepsy, moderate to severe developmental delay, autistic features, structural brain abnormalities, hypotonia, short stature and variable renal and cardiac abnormalities. HnRNP U is essential for mammalian development as lethality results by embryonic day 11.5 in mice carrying homozygous hypomorphic mutations^26^. Furthermore, homozygous pathogenic mutations have yet to be reported in humans^27^. Conditional loss of *Hnrnpu* in mouse cardiomyocytes was also associated with a lethal dilated cardiomyopathy and widespread transcriptional and splicing dysregulation including known cardiomyopathy disease genes^16^. However, the transcriptomic and physiologic effects of *Hnrnpu* haploinsufficiency in the brain have yet to be characterized.

Here we assess the neurophysiological consequences and face validity of an *Hnrnpu* mouse disease model using *in vivo* developmental, morphological, electrophysiological, and behavioral studies. We also perform a comprehensive cell-type- and brain region-specific characterization of reduced *Hnrnpu* levels on gene expression using scRNAseq at a single postnatal time point. Using these data, we generate cell-type-specific disease expression signatures and identify vulnerable cell types in the mutant mouse brain. We then compare these signatures to publicly-available gene expression signatures of cells treated with small molecules in order to identify compounds that correct disease-associated transcriptomic changes towards a normal state. Overall, this work provides a framework for high-resolution phenotyping of models of transcriptome-mediated diseases and outlines important considerations for the future development of targeted therapies for *HNRNPU* DEE.

## Results

### Generation of an *Hnrnpu* knockout mouse model

HnRNP U expression is widespread in the brain, yet particularly concentrated within the cerebellum, hippocampus and neocortex^28^. In mouse primary cell cultures derived from the neocortex and hippocampus, hnRNP U co-stained with the neuronal marker Map2, and markers of neuronal subtypes including inhibitory neurons (Gad67), as well as cortical pyramidal neurons of deep (Ctip2) and superficial (Satb2) lamina **(Supplemental Figure 1A-D, Supplemental Figure 2A-D)**. HnRNP U also co-stained with the astrocyte marker Gfap **(Supplemental Figure 1E, Supplemental Figure 2E)**. For all cells examined, hnRNP U expression appeared confined to the nucleus, as previously reported^7,8^.

Most pathogenic mutations in *HNRNPU* are loss-of-function **(Figure 1A, Supplemental Table 1)**. We therefore targeted exon 1 of mouse *Hnrnpu—*a region in which at least five nonsense mutations were identified in human patients—to induce a constitutive out-of-frame deletion **(Figure 1A-B, Supplemental Table 1)**. A founder containing a heterozygous 113-bp deletion (herein referred to as *Hnrnpu*^*+/*113DEL^ or HET) with resulting premature stop codon in exon 2 was identified and used to expand the line, which was subsequently maintained on a C57BL/6NJ background **(Figure 1B, Supplemental Figure 3A)**.

**Figure 1.**
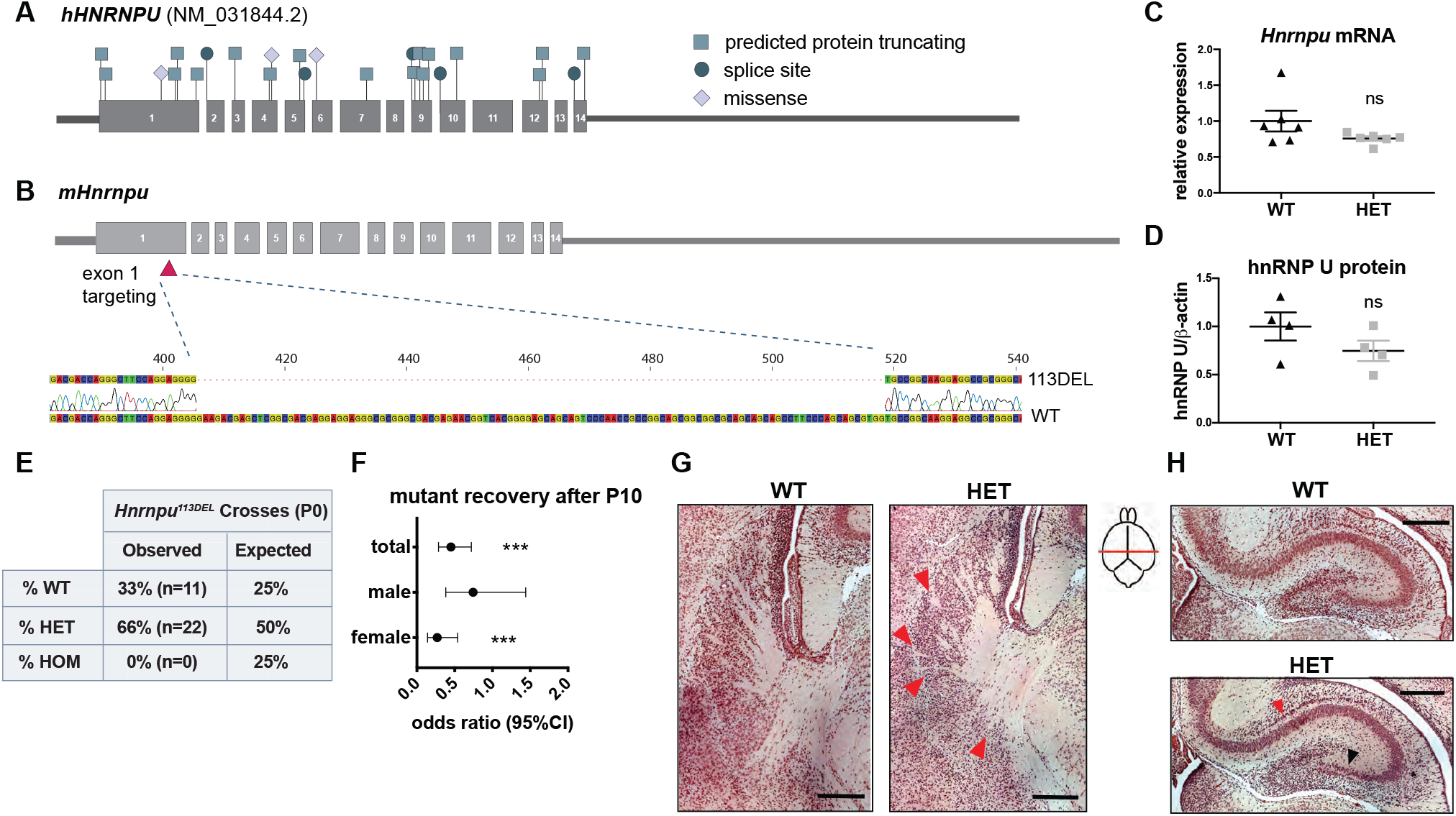
Generation of a mouse model of *HNRNPU* haploinsufficiency. **(A)** Location of all predicted pathogenic variants, including protein truncating, splice site and missense, reported in the literature across all 14 coding exons of human *HNRNPU* (NM_031844.2). **(B)** Location of the CRISPR-induced 113-bp deletion in exon 1 of mouse *Hnrnpu*. **(C)** *Hnrnpu* expression relative to *Cyc1* reference gene. Welch’s t-test p= 0.15 (two-tailed, t= 1.65, df= 5). Error bars= SEM. **(D)** Mouse hnRNP U protein expression quantified by densitometry and normalized to mouse β-actin (n= 4 P0 cortices per genotype). Unpaired t-test p= 0.21 (two-tailed, t=1.4, df=6). Error bars= SEM **(E)** Observed versus expected Mendelian ratios from *Hnrnpu*^+/113DEL^ intercrosses (n= 10 litters). **(F)** Odds (with 95% confidence interval) of recovering HET pups beyond P10 compared to P0. Fishers exact test p-values: females= 2×10^−4^, males= 0.41, total= 8×10^−4^. **(G)** Representative images of WT and HET striatum. Red arrowheads indicate region with fewer axon fascicles. **(H)** Representative images of WT and HET hippocampus. The red and black arrowheads indicate an abnormal layer of cells in *Hnrnpu*^+/113DEL^ CA1 and CA3, respectively (n= 3 animal per genotype). Scale bar = 250 μm.

Evaluation of both mRNA and protein levels obtained from cerebral cortex showed only a partial 20-25% reduction of *Hnrnpu* expression in *Hnrnpu*^+/113DEL^ mice compared to the expected 50% **(Figure 1C-D, Supplemental Figure 3B)**. Furthermore, we did not detect a truncated form of hnNRP U when using an antibody that binds N-terminal to the deletion breakpoint **(Supplemental Figure 3C)**. We compared this modest decrease in mutant *Hnrnpu* expression to two isogenic human induced pluripotent stem cell (hiPSC)-derived neuronal models containing distinct heterozygous loss-of-function mutations in exon 2 of *HNRNPU* (HET1= 1bp duplication and HET2= 10bp deletion). Gene expression was assessed at day 10 and 45-47 of human cortical organoid (hCO) induction and compared to an isogenic WT control and a non-isogenic WT line serving as an independent control. While both time points showed a tendency towards reduction in *HNRNPU* expression, the degree of reduction for both mutants varied based on the loading control used, with *ATP5F1* showing a more pronounced decrease ranging from 38-76% (**Supplemental Figure 4A-B**) compared to 17-39% for GAPDH (**Supplemental Figure 4C-D**). In line with our mouse expression data, these findings highlight the inherent variability in *HNRNPU* gene expression and suggest the presence of compensatory mechanisms in response to protein-truncating variants in *HNRNPU*.

*Hnrnpu*^+/113DEL^ intercrosses did not produce any viable homozygous mutant progeny, consistent with embryonic lethality **(Figure 1E)**. *Hnrnpu*^+/113DEL^ mice also demonstrated increased perinatal mortality; compared to postnatal day 0 (P0), half of the mutants were recovered on or beyond P10 (Fisher’s exact test (FET) p= 8×10^−4^) **(Figure 1F)**. This difference appeared to be largely driven by a disproportionately higher loss of female mutants, with 0.27 odds of recovering female mutants (FET p= 2×10^−4^, 95% CI) compared to 0.74 for males (FET p= 0.41, 95% CI) **(Figure 1F)**.

### Reduced hnRNP U causes abnormal brain development

Patients with *HNRNPU* mutations present with a variety of central nervous system abnormalities, including neuronal migration defects, enlarged lateral ventricles, corpus callosum defects, delayed myelination and mild holoprosencephaly^21–23^. We did not observe differences in *Hnrnpu*^+/113DEL^ brain size and corpus callosum morphology compared to wildtype (WT) at P0 **(Supplemental Figure 5A)**. Further examination of *Hnrnpu*^+/113DEL^ brains shows no significant changes in cortical thickness and hippocampal width **(Supplemental Figure 5B)**. However, *Hnrnpu*^+/113DEL^ mice showed fewer axon fascicles in the striatum **(Figure 1G)**. Unlike WT, these fascicles appeared spotty and fragmented (red arrowheads, **Figure 1G)**, suggesting a change in projection trajectory. Furthermore, in the *Hnrnpu*^+/113DEL^ hippocampus, cells in CA1 (red arrowhead) and CA3 (black arrowhead) were abnormally organized into two distinct layers indicating a lamination defect **(Figure 1H)**.

#### *Hnrnpu*^+/113DEL^ pups show global developmental delay

Patients with *HNRNPU*-associated DEE frequently demonstrate axial hypotonia along with moderate to severe developmental delay, primarily manifesting as delayed motor skills and severe speech impairment^21–23^. We therefore evaluated early physical and sensorimotor development including growth, righting reflex, negative geotaxis and vertical screen hold in the first two weeks of life.

At birth, mutant pups weighed on average 10% less than WT controls (MWU, permuted p= 5×10^−3^) **(Supplemental Figure 6A)**. This growth impairment was further exacerbated throughout the postnatal period, with P12 mutants weighing roughly 25% less than controls (MWU permuted p< 0.01 for all time points) **(Figure 2A)**. This degree of growth impairment persisted throughout the juvenile period into adulthood (MWU permuted p< 0.01 for all time points) **(Supplemental Figure 6B)**.

Despite weighing significantly less, *Hnrnpu*^+/113DEL^ pups showed a subtle increase in latency to fall at P6 in the vertical screen test **(Supplemental Figure 6C)**. Mutants also showed a modest impairment in both righting reflex at P10 and the 90° negative geotaxis (the time it takes to right 90 degrees from a downward facing position on a wired mesh at a 45° angle) at P12, highlighting a trend towards delayed sensorimotor function (MWU permuted p= 1×10^−3^ and p= 4×10^−3^, respectively) **(Supplemental Figure 6D-E)**. There was no significant difference in 180° negative geotaxis for any of the time points evaluated **(Supplemental Figure 6F)**.

To further assess developmental delay in *Hnrnpu*^+/113DEL^ pups, we evaluated separation-induced ultrasonic vocalizations (USVs). USVs are functionally important signals that elicit maternal retrieval and care^29^. Deficits in pup USVs have been reported in various rodent models of neurodevelopmental disease, most notably in models of human communication disorders such as autism and verbal dyspraxia^30–35^. Evaluation of USVs from WT mice revealed the canonical inverted-U shape from P3 to P11, characteristic of normal pups^36^ **(Figure 2C)**. The number of WT pup calls increased steadily and peaked at P7 **(Figure 2C)**. Conversely, *Hnrnpu*^+/113DEL^ pups showed clear deficits in USVs **(Figure 2B)**, including a striking reduction in the number of calls, particularly at P5 and P7 (MWU permuted p= 7×10^3^ and < 1×10^−4^, respectively), with an atypical trajectory characterized by a slow increase in the number of calls that peaked around P9 **(Figure 2C)**. Further analysis of USV acoustic properties of mutants at P5 and P9 revealed a shorter duration (MWU permuted p= 2×10^−3^ and < 1×10^−4^, respectively) and overall higher frequency (MWU permuted p= 7×10^−3^ and 0.02, respectively) compared to control calls **(Figure 2D, E)**. Moreover, mutant vocalizations also trended towards an increased peak amplitude, although this observation was only significant at P9 (MWU permuted p= 3×10^−3^) **(Figure 2F)**. Overall, these data, combined with growth and milestone studies, are consistent with global developmental delay in *Hnrnpu*^+/113DEL^ mice.

**Figure 2.**
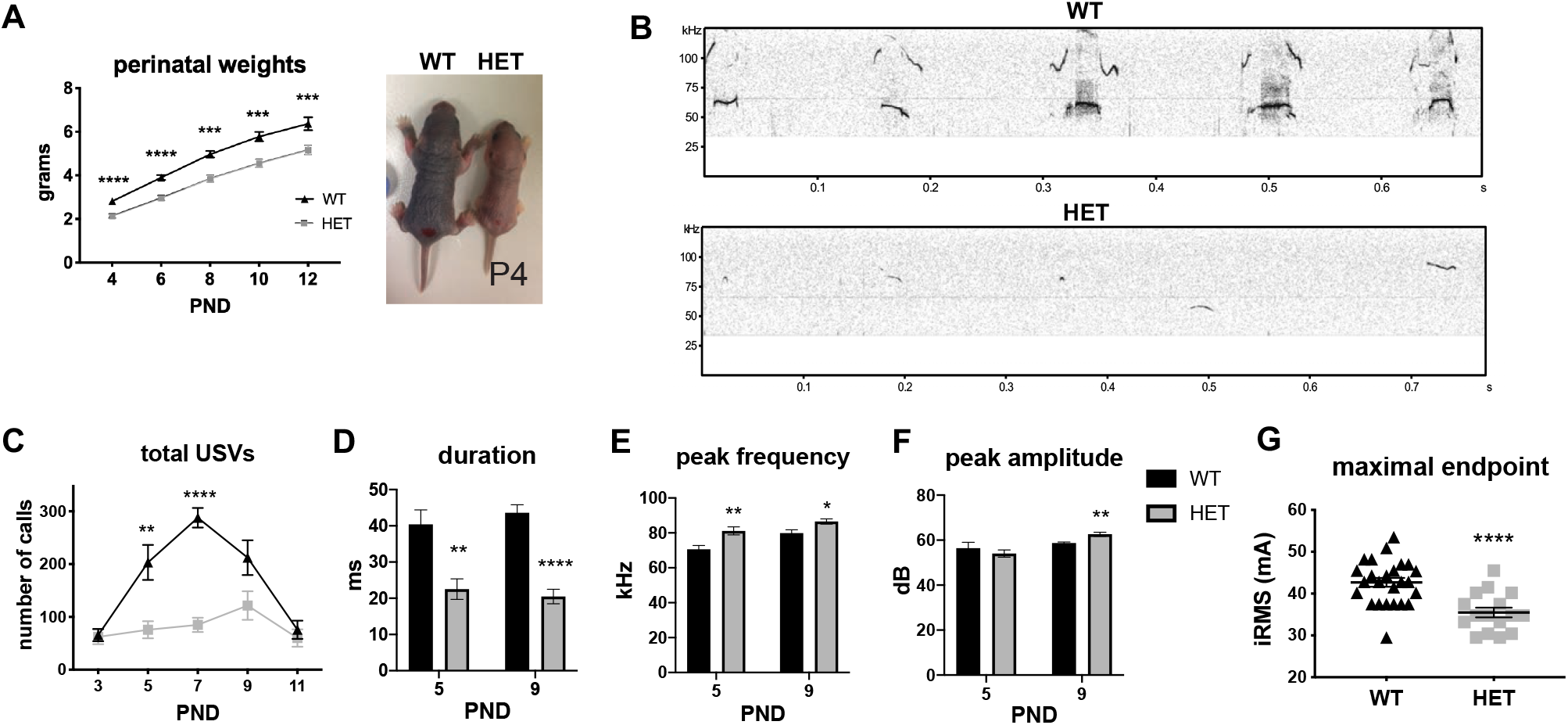
*Hnrnpu*^+/113DEL^ mouse phenotypes. **(A)** Body weights obtained in the perinatal period (n= 16 per genotype). Permuted Mann-Whitney U (MWU) p-values: P4 and P6< 1×10^−4^, P8= 2×10^−4^, P10= 1×10^−4^, P12= 1.5×10^−3^. Representative image of a WT and HET P4 pup. **(B)** Representative image of ultrasonic vocalization spectrograph. **(C)** Quantity of pup calls over a 3 min time interval (n= 17 WT, 16 HET pups). Permuted MWU p-values: P3= 0.61, P5= 6.8×10^−3^, P7< 1×10^−4^, P9= 0.06, P11= 0.49. **(D)** Average pup call duration. Permuted MWU p-values: P5= 2.1×10^−3^, P9< 1×10^−4^. **(E)** Average peak frequency (i.e. pitch). Permuted MWU p-values: P5= 6.9×10^−3^, P9= 0.02. **(F)** Average peak amplitude (i.e. loudness). Permuted MWU p-values: P5= 0.43, P9= 3.3×10^−3^. For all qualitative USV analyses, n= 10 for each genotype at P5, n=11 for each genotype at P9. **(G)** Maximal seizure ECT endpoint (n= 25 WT, 16 HET adults). MWU p< 1×10^−4^. iRMS= root mean square current. PND= postnatal day. Error bars= SEM.

#### *Hnrnpu*^+/113DEL^ adults exhibit seizure susceptibility

Given the presence of early-onset seizures in patients with *HNRNPU* mutations^21–23^, we assessed *in vivo* spontaneous and evoked excitability phenotypes using electroencephalography (EEG) and electroconvulsive threshold (ECT) studies, respectively. Despite over 300 total hours of video EEG recordings among seven *Hnrnpu*^+/113DEL^ adults, there was no evidence of spontaneous generalized epileptiform activity **(Supplemental Figure 7A)**. Moreover, no spontaneous seizure-like behaviors or sudden death were observed following routine handling of this mouse line at any age. However, mutants demonstrated a significantly lower threshold for maximal tonic hindlimb extension seizures, consistent with a greater seizure predisposition (MWU p< 1×10^−4^) **(Figure 2G)**.

In light of the moderate to severe intellectual disability, along with motor and neuromuscular impairments observed in patients with *HNRNPU* mutations^21–23^, adult mice were surveyed for exploratory activity, gait, and learning and memory. Results revealed only modest differences between *Hnrnpu*^+/113DEL^ and WT adult mice **(Supplemental Figure 7B-P)**.

### Single-cell RNA-sequencing of the neocortex and hippocampus reveals ubiquitous *Hnrnpu* expression

We performed scRNAseq on neocortical and hippocampal samples obtained from *Hnrnpu*^+/113DEL^ and WT littermates at a single postnatal time point (P0). For both brain regions, we evaluated two pups of each genotype, including one of each sex. Cortices and hippocampi were dissected from different mice originating from separate litters. In total, we sequenced 18,171 neocortical cells and 21,487 hippocampal cells.

Using Seurat^37,38^, we harmonized expression data across WT and *Hnrnpu*^+/113DEL^ cells before performing unsupervised clustering **(Supplemental Figure 8A, B)**. We then combined cell clusters into major cell classes based on expression of well-established canonical marker genes **(Figure 3B, E; Supplemental Table 2)**^39,40^. In total, we identified 13 distinct cell populations for each brain region **(Figure 3A, D)**. Overlapping cell populations included proliferative cells, radial glia, intermediate progenitors, oligodendrocyte precursor cells, and inhibitory subpopulations, including SST and VIP positive interneurons **(Figure 3A, D)**. In the neocortex, we identified additional inhibitory neuron clusters, including LGE-derived interneurons, interneuron progenitors, and spinal projection neurons **(Figure 3D)**. We classified neocortical pyramidal neurons into two major populations: upper layers 2 through 4 and deeper layers 5 and 6. We classified hippocampal pyramidal neurons based on their respective hippocampal subfield, including the dentate gyrus, CA1, CA2 and CA3, subiculum, and entorhinal cortex **(Figure 3A)**. *Hnrnpu* was expressed ubiquitously across all neocortical and hippocampal cell populations, though was slightly increased in proliferative cells (i.e. neural stem cells) **(Figure 3C, F)**.

**Figure 3.**
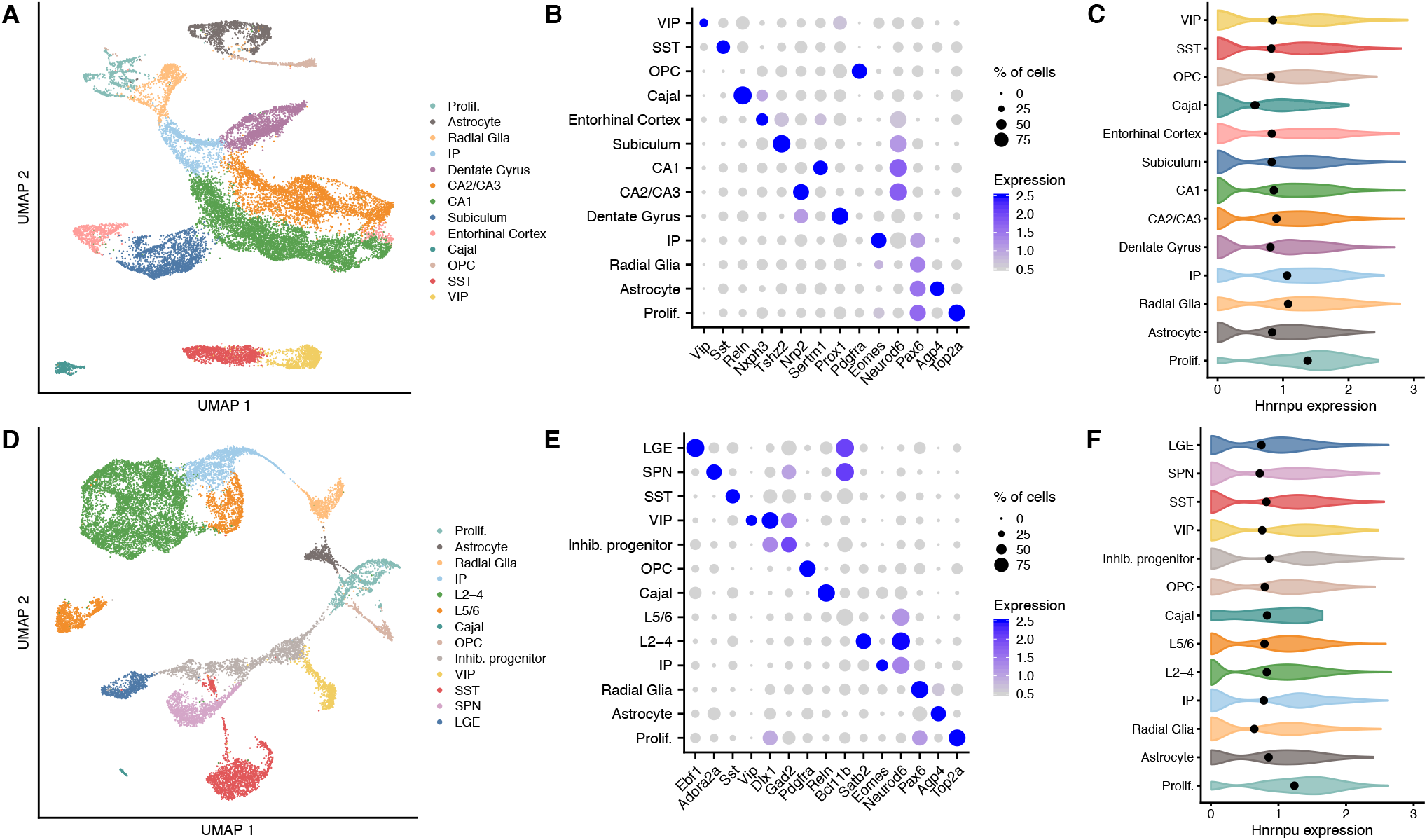
Single-cell RNA-sequencing of wildtype and mutant neocortical and hippocampal cells. **(A)** UMAP plot of all hippocampal cells, colored by cell type. Prolif: proliferative cells; IP: intermediate progenitors; Cajal: Cajal-Retzius cells; OPC: oligodendrocyte precursor cells; SST: SST+ interneurons; VIP: VIP+ interneurons. **(B)** Expression of canonical cell type markers used for annotating each hippocampal cell population. **(C)** Expression of *Hnrnpu* in each population of WT hippocampal cells. **(D)** UMAP representation of all neocortical cells, colored by cell type. L2-4: upper layer (layers 2-4) pyramidal neurons; L5/6 deep layer (layers 5 and 6) pyramidal neurons. **(E)** Expression of canonical cell type markers used for annotating each neocortical cell population. **(F)** Expression of *Hnrnpu* in each population of WT neocortical cells.

### Cell-type-specific differential gene expression analysis

We next performed differential gene expression to identify cell-type-specific perturbations in the *Hnrnpu*^+/113DEL^ brain. We compared gene expression profiles from mutant and WT cells of each population using a linear mixed model **(Figure 4A, C)**. In the hippocampus, we detected 955 differential expression events (FDR q< 0.05 and expression change > 10%), composed of 679 unique differentially expressed genes (DEGs) **(Supplemental Table 3)**. In the neocortex, we detected 454 differential expression events, composed of 303 unique DEGs **(Supplemental Table 4)**. Notably, in the hippocampus there were substantially more downregulated differential expression events (698 genes; 73%) than upregulated (257; 27%). This pattern was not as evident in the neocortex, in which 230 differential expression events resulted in downregulation (51%) compared to 224 that resulted in upregulation (49%). Effect sizes were generally modest. The average absolute log fold change was 0.24 among hippocampal differential expression events and 0.25 among neocortical differential expression events.

### Downregulated genes converge on neuronal processes and pathways

In effort to determine whether DEGs were enriched for certain biological pathways, we performed gene ontology analyses. Downregulated genes in both the mutant hippocampus and neocortex were enriched for several ontologies relevant to the disease phenotype, including neuron projection development, axon guidance, neuron migration, and glutamatergic synaptic signaling (**Figure 4E, F**). These dysregulated pathways—namely axon guidance and neuronal migration— are supported by the morphological phenotypes observed in the mutant hippocampus and striatum (**Figure 1F, G**) **(Supplemental Table 5)**. Meanwhile, upregulated genes were more strongly enriched for ontologies relevant to cellular growth, differentiation, protein translation and localization **(Supplemental Table 5)**.

**Figure 4.**
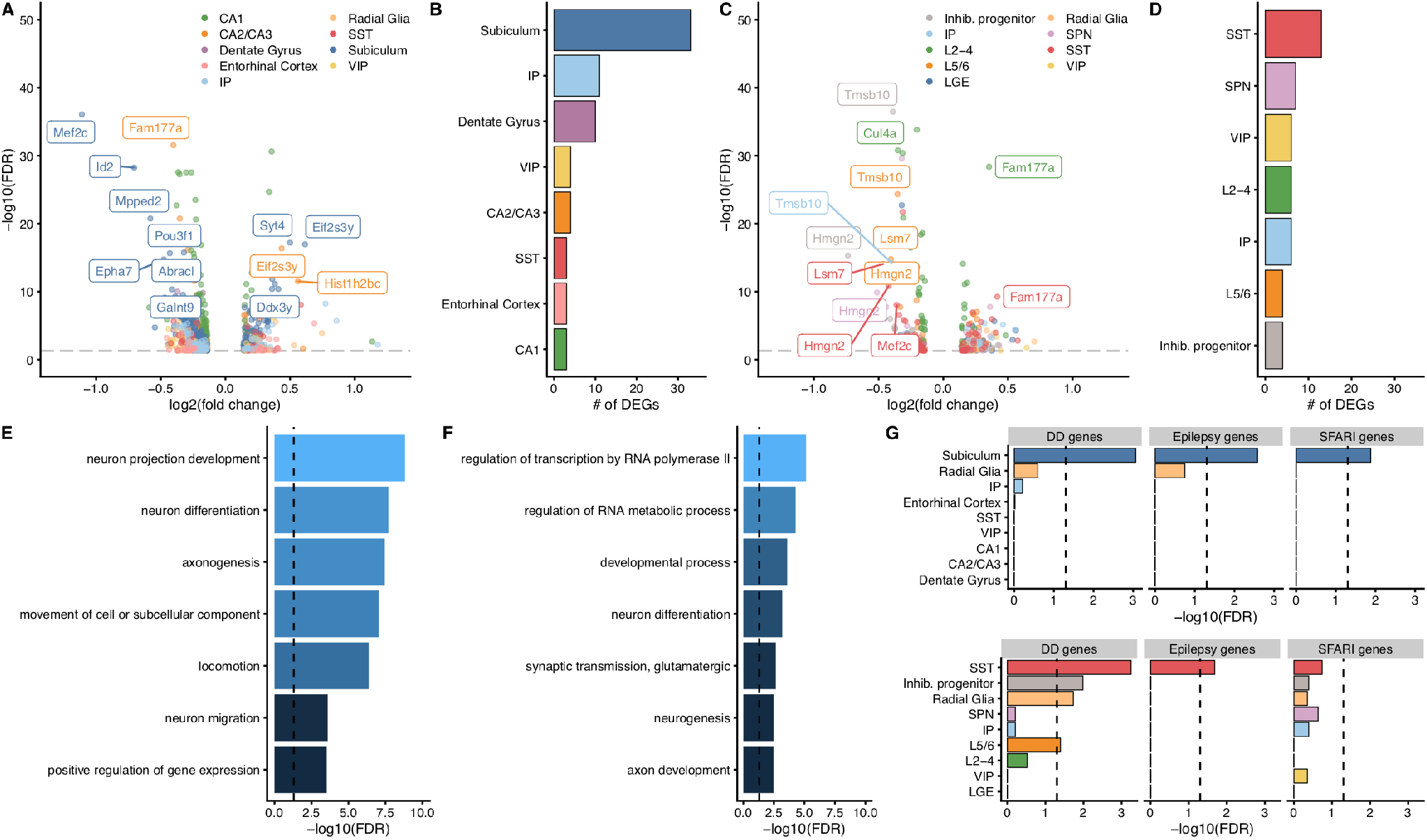
Cell-type-specific dysregulation of gene expression in the hippocampus and neocortex. **(A)** Volcano plot representing differentially expressed genes in the hippocampus. **(B)** Burden of differentially expressed genes in each hippocampal cell type based on downsampled data. **(C)** Volcano plot representing differentially expressed genes in the cortex. **(D)** Burden of differentially expressed genes in neocortical cell types based on downsampled data. **(E, F)** Gene ontology analysis of downregulated genes in the hippocampus and neocortex, respectively. **(G)** Enrichment of developmental delay (DD), epilepsy, and autism (SFARI) genes among nominally significant DEGs in hippocampal (upper panel) and neocortical (lower panel) cell types.

### An increased burden of DEGs in excitatory neurons of the mutant subiculum

We next tested whether certain cell types are more vulnerable to the reduced expression of *Hnrnpu* than others. First, we downsampled the data to compare the same number of cells across all cell types (n= 300 cells per cell type; **Supplemental Figure 9A,B**). We re-ran the differential gene expression analysis on these downsampled data and found that excitatory neurons derived from the subiculum showed the highest burden of DEGs across all cortical and hippocampal cell types **(Figure 4B, D)**. The next strongest burden was identified in SST+ interneurons of the neocortex. Interestingly, the neocortical SST+ interneurons had more DEGs than hippocampal SST+ interneurons.

Because patients with *HNRNPU* haploinsufficiency have autistic features, developmental delay, and seizures, we next tested for the enrichment of genes associated with these conditions among each population of cells. Here, we relaxed the significance threshold for DEGs to an FDR < 0.1 and expression change of at least 5% given the relatively small sizes of these gene sets and weak expression of these disease genes at P0. Strikingly, we observed an overrepresentation of developmental delay, epilepsy, and autism genes among the downregulated genes in the subiculum-derived pyramidal cells **(Figure 4G)**. No other cell type showed this strong of an enrichment for all three disease gene sets. We also observed an enrichment of developmental delay and epilepsy genes among downregulated genes in SST+ neocortical interneurons **(Figure 4G)**. Altogether, these results point to the subiculum as a potentially vulnerable region that may play an important role in mediating the pathophysiology underlying *HNRNPU-*DEE.

### *Mef2c* is the most downregulated gene in the mutant subiculum

The most downregulated gene observed in the subiculum was *Mef2c*, which showed a roughly 50% decrease in expression (log_2_ fold change = -1.11, FDR = 8 × 10^−37^) **(Figure 5A)**. This effect size was among the largest observed fold-changes of genes differentially expressed in both the neocortex and hippocampus. Expression of *Mef2c* in the P0 hippocampus is primarily confined to both subiculum-derived pyramidal neurons and SST+ interneurons **(Figure 5A)**. Its expression is more widespread in the P0 cortex, including expression in upper and deep layer pyramidal neurons, along with SST+ and VIP+ interneurons **(Figure 5B)**. Interestingly, despite this widespread expression, the only other cells in which *Mef2c* was significantly downregulated were SST+ interneurons, yet to a lesser degree than subiculum-derived pyramidal neurons (FDR q= 9 × 10^−9^; log_2_ fold-change = -0.36) **(Figure 5A, B)**. This finding further highlights the presence of cell-type-specific effects upon the loss of ubiquitously expressed hnRNP U protein.

**Figure 5.**
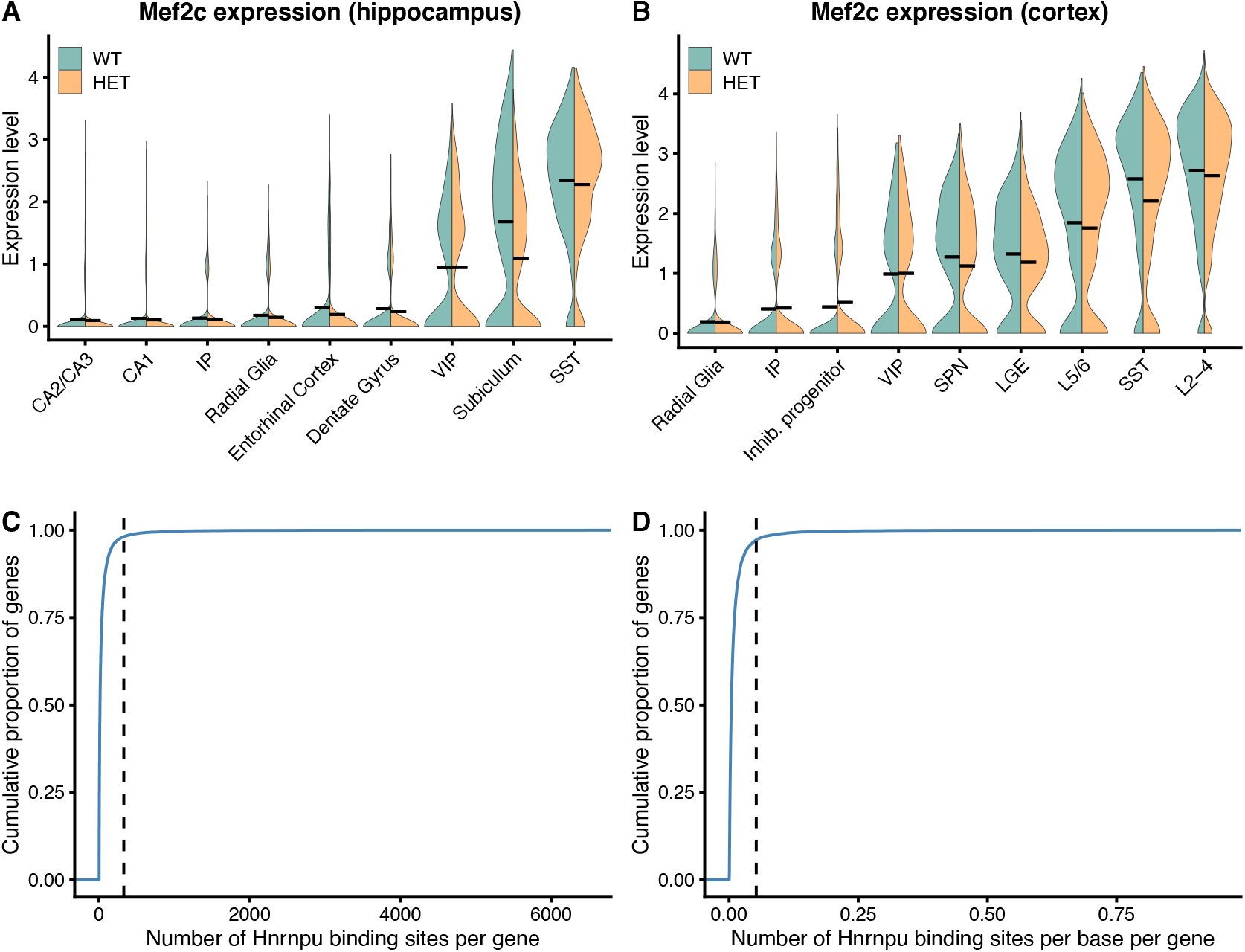
Dysregulation of *Mef2c*. **(A, B)** Expression of *Mef2c* in wildtype and mutant hippocampal and cortical cells. Black lines represent median expression levels in each genotype. **(C)** Cumulative distribution plot representing Hnrnpu binding sites per gene, derived from CLIP-seq data performed on cardiac tissue. Dotted line represents the number of Hnrnpu binding sites in *Mef2c*. **(D)** Same as (C), except that the number of binding sites is normalized by transcript length.

To assess whether hnRNP U likely functions to directly regulate the expression of *Mef2c*, we examined the number of hnRNP U binding sites for each gene expressed in the brain of mice. Using available hnRNP U CLIP-sequencing data derived from mouse cardiac tissue, we found that *Mef2c* contains more hnRNP U binding sites than 99% of the other genes assessed **(Figure 5C)**. This effect persisted when we normalized the number of binding sites by gene length **(Figure 5D)**. Together, these results suggest that *Hnrnpu* plays an important role in regulating expression of *Mef2c*, at least at this particular timepoint.

### Candidate compounds for transcriptomic reversal of the *Hnrnpu*^*+/113DEL*^ disease expression signature

Because heterozygous loss of *Hnrnpu* leads to widespread cell-type-specific dysregulation of gene expression, identifying targeted therapeutics could prove especially challenging. However, transcriptomic signature reversal—a paradigm well-developed in cancer— may provide one particularly promising avenue for drug discovery for both this disease and other neurodevelopmental diseases caused by genes that directly influence the transcriptome. This paradigm posits that if gene expression changes underlie the pathophysiology of a particular disease, then correction of this transcriptomic signature toward a normal state may have therapeutic potential. Transcriptomic reversal requires the comparison of a disease gene expression signature and the gene expression signatures of cells treated with small molecules. Small molecules that elicit expression changes most anticorrelated with the disease signature are prioritized for further validation. The Connectivity Map (CMAP)^44,45^ provides publicly available expression signatures derived from cancer cell lines treated with thousands of small molecules.

Transcriptomic reversal approaches that have leveraged the CMAP and other resources have not only successfully identified targeted therapeutics in cancer^46–48^, but for other diseases too, including diabetes and inflammatory bowel disease^49,50^. However, this approach has not been successfully applied to transcriptome-mediated neurodevelopmental conditions. Given the pleiotropic, cell-type-specific effects of *Hnrnpu* haploinsufficiency in disease-relevant brain regions, we expect that this approach will require scRNAseq-derived signatures.

To examine the importance of cell-type-specific effects, we compared compounds predicted to reverse the subiculum gene expression signature versus compounds predicted to reverse a “pseudo-bulk” hippocampal signature (i.e. the average gene expression changes across all cell types). We specifically focused on the downregulated genes in the disease signatures, as these genes were enriched for biologically relevant pathways and disease genes. The CMAP uses a “connectivity score” to assess each compound’s ability to reverse the query signature^44^. This score ranges from -100 to +100, with a score of -100 indicating complete reversal.

For the subiculum-derived signature, 128 compounds received a Connectivity Score less than the CMAP’s recommended cutoff of -90 **(Supplemental Figure 10A, Supplemental Table 6)**. 98 compounds received a score below this threshold for the pseudo-bulk derived signature **(Supplemental Figure 10B, Supplemental Table 6)**. Only 40 (31%) of these candidate compounds overlapped between the two queries **(Figure 6E)**. Furthermore, among the top 20 compounds prioritized per signature, only four compounds overlapped (linifanib, Merck60, etinostat, and BMS-345541) **(Supplemental Figure 10C, D)**. The classes of compounds among these 20 compounds prioritized for each signature were also different. An overwhelming majority of pseudo-bulk prioritized compounds were HDAC inhibitors (70%), whereas the most common drug class for the subiculum-prioritized signatures were tubulin inhibitors and microtubule stabilizing agents (25%).

Therefore, if transcriptional dysregulation of a specific cell type, such as excitatory neurons in the subiculum, contributes substantially to the pathophysiology underlying *HNRNPU* haploinsufficiency, bulk RNA-sequencing may not generate adequate signatures to identify the compounds most likely to target the relevant disease mechanisms. Experimental validation will be required to verify that these candidate compounds in fact reverse the transcriptomic signatures and rescue Hnrnpu disease phenotypes. Nonetheless, these results highlight the potential importance of deriving cell-type-specific disease expression signatures for transcriptomic signature reversal approaches.

## Discussion

Identifying and modeling germline mutations associated with developmental and epileptic encephalopathies provides the unique opportunity to develop targeted therapeutics^51^. While the majority of these mutations tend to occur in genes that encode ion channels or synaptic transmission proteins, a subset of these genes can be thought of as causing disease through their effects on the transcriptome^52,53^. Unfortunately, elucidating disease mechanisms for this subset of genes is often difficult, as the encoded proteins regulate the expression of thousands of target genes. To address this problem, using our precision genetic mouse model, we evaluated the potential utility of a transcriptome-guided precision medicine approach for hnRNP U-mediated neurodevelopmental disease, relying on brain-region and single cell-level gene expression profiles to highlight vulnerable cell types and key dysregulated genes.

### Near wildtype expression of hnRNP U is necessary for normal development

Intriguingly, we observed that loss of a single copy of mouse *Hnrnpu* results in a 20-25% reduction in expression, instead of a 50% loss as predicted by the protein truncating nature of the frameshift mutation. Despite this partial reduction in expression, heterozygous intercrosses failed to produce any homozygous progeny, consistent with embryonic lethality in mice. These findings are also consistent with the observation that mutant ES cells containing embryonically-lethal homozygous hypomorphic *Hnrnpu* mutations demonstrate only a 40-80% decrease in *Hnrnpu* transcript levels^26^. Furthermore, WT and *Hnrnpu*^+/113DEL^ crosses demonstrated skewed Mendelian ratios with greater postnatal loss of female mutants. This may be caused by compromised X-chromosome inactivation in these mutant female mice, due to reduced *Xist* RNA coating mediated by hnRNP U^54^. For comparison, we also evaluated *HNRNPU* expression in a human disease model, and found varying degrees of *HNRNPU* transcript reduction in induced pluripotent stem cells and cortical organoids, ranging from a partial decrease similar to that observed in mice, to over 50% reduction.

While the mechanism underlying this partial loss of *Hnrnpu* expression remains unknown, potential explanations include the enhanced stability of *Hnrnpu* transcripts, such as through autoregulation via self 3’UTR binding^15^ or the compensatory upregulation of *Hnrnpu* expression through mechanisms like transcriptional adaptation^55^. Differentiating between these possibilities could unveil future avenues for molecular therapy, such as targeting the transcriptional adaptation mechanism to further upregulate *Hnrnpu* expression.

### *Hnrnpu*^+/113DEL^ mice model aspects of *HNRNPU* developmental and epileptic encephalopathy

Here we performed the first neurophysiological characterization of an *HNRNPU* neurodevelopmental disease model. *Hnrnpu*^+/113DEL^ mice displayed several phenotypes that overlap in presentation with the human disorder^21–23^. A general morphological evaluation of the brain revealed the presence of both axon pathfinding and hippocampal lamination defects. These findings were paralleled by global developmental delay as evidenced by impaired growth, delayed sensorimotor function and striking deficits in separation-induced pup ultrasonic vocalizations. The short and high-pitched presentation of mutant calls are likely less effective as communicative signals^56,57^, and may contribute to the exacerbation of the physical growth impairment noted during the perinatal period from poor maternal care. Mutant mice that survive to adulthood also exhibited greater seizure susceptibility. These findings support the utility of this model to further investigate and interrogate the molecular underpinnings of *HNRNPU*-mediated disease phenotypes. Our study also builds upon prior behavioral work conducted in an independent heterozygous *Hnrnpu* knockout model that demonstrated altered circadian-mediated locomotor and metabolic activity in mice carrying an out-of-frame deletion of exons 3 through 6^58^.

### HnRNP U is important for sustaining expression of neuronally-expressed genes

Through single-cell RNA-sequencing of the neocortex and hippocampus, we identified widespread transcriptional dysregulation across all the neuronal cell types examined in mutant mice. The overall magnitude of these expression changes was generally modest—a phenomenon similar to what has been observed in Rett Syndrome, both in post-mortem human tissue and *Mecp2* mouse models, as well as a mouse model of CHD8-mediated neurodevelopmental syndrome^59–61^. Nevertheless, this modest transcriptional dysregulation could also result from mouse compensatory or other species-specific differences from the human condition. To better understand the significance of these results, it is imperative to also evaluate and compare to the transcriptional state of humanized *HNRNPU* disease models such as iPSC-derived neurons or brain organoids.

In both the neocortex and hippocampus, we observed downregulation of genes that were particularly enriched for important, disease-relevant ontologies, including neuronal migration and axon guidance pathways—findings which were supported by the morphological defects observed in the *Hnrnpu*^+/113DEL^ mouse striatum along with hippocampal CA1 and CA3 subfields. These results suggest an important role for hnRNP U in sustaining the expression of many disease-relevant, neuronally-expressed genes.

### Cell-type-specific effects upon heterozygous loss of *Hnrnpu*

Interestingly, we found that the greatest burden of gene expression dysregulation upon heterozygous loss of *Hnrnpu* occurred in pyramidal cells of the subiculum. Downregulated genes in this cell type were also enriched for known neurodevelopmental disease genes including those associated with autism, developmental delay and epilepsy. Although the subiculum functions as the primary output of the hippocampus and is important for normal hippocampal-related functions, such as learning and memory and stress response, this brain region has also been implicated in pathological conditions, such as the generation and spread of temporal lobe seizures^62–66^. While the location of seizure onset was not reported for most individuals with *HNRNPU* mutations, two patients reportedly had temporal lobe epilepsy^21,23^. Furthermore, haploinsufficiency of the most downregulated gene in the subiculum, Mef2c, is associated with a neurodevelopmental disorder showing significant phenotypic overlap to patients with *HNRNPU* mutations^67,68^. While these findings only reflect a single snapshot of time and do not account for other transcriptional insults that may have incurred during development and adulthood, these results do demonstrate relevant cell-type and brain-region specific gene expression changes that may contribute to the general dysfunction and subsequent pathological phenotypes.

### Transcriptomic reversal of mutant cell-type-specific signatures as a therapeutic approach

Widespread and cell-type-specific transcriptomic defects present a challenge with regard to pinpointing targeted therapeutics. However, transcriptomic signature reversal may provide one route forward. This approach has been successful in identifying compounds that ameliorate seizures in non-genetic models of epilepsy^70^. Through querying the Connectivity Map^44,45^, we find numerous compounds that may reverse disease gene expression signatures. We find that the prioritized compounds, however, are sensitive to the use of cell-type-specific versus bulk-derived expression signatures. Therefore, identifying the most vulnerable cell types for each transcriptome-mediated epilepsy gene may be at least one crucial factor for the successful application of this approach in neurodevelopmental diseases.

### Conclusion

Here we characterized the neurophysiological and cell-type-specific transcriptomic effects of a mouse model of *HNRNPU* developmental and epileptic encephalopathy. Although a heterozygous loss-of-function mutation resulted in only 20-25% loss of *Hnrnpu* expression, mutant mice demonstrated abnormal neurodevelopment and other features reminiscent of the human disorder. We observed pervasive gene expression changes of modest effect that occurred in a cell-type-specific manner, including many neuronally-expressed genes that converge on important neuronal pathways. This was accompanied by a nearly 50% reduction in expression of a single neurodevelopmental disease gene, *Mef2c*, in pyramidal neurons of the subiculum—a cell-type particularly vulnerable to heterozygous loss of *Hnrnpu*. These results imply a complex role for hnRNP U in gene expression regulation—a notion supported by its known function in regulating multiple levels of gene expression. This complexity is likely further amplified by many factors including context-dependent interactions with other gene expression regulators, developmental timing and species-specific variation that were not examined as part of this work but warrant further study. These results also support the exploration of alternative therapeutic approaches, with transcriptomic signature reversal of key, vulnerable cell-types as a promising, novel strategy for transcriptome-mediated neurodevelopmental disease.

## Materials and Methods

### Mouse husbandry

*Hnrnpu*^+/113DEL^ mice were generated through The Jackson Laboratory’s Genome Engineering Technology core using a CRISPR-Cas9 strategy targeting exon 1 of mouse *Hnrnpu*. Of 15 founder mice that survived, 6 appeared most promising based on TIDE analysis and were further evaluated using TOPO-TA cloning to validate the corresponding mutant alleles. A founder containing an out-of-frame 113-bp deletion was further expanded and maintained on a C57BL/6NJ background. All experiments were performed on the inbred background except for ECT studies, which were performed on the F1 hybrid background C57BL/6NJ (005304 JAX stock) x FVB/NJ (001800 JAX stock), as mutants on an inbred C57BL/6NJ were significantly smaller than WT controls. WT littermates were used as controls in all experiments. All mice were maintained in ventilated cages with controlled humidity at ∼60%, 12h:12h light:dark cycles (lights on at 7:00AM, off 7:00PM) and controlled temperature of 22–23°C. Mice had access to regular chow and water, ad libitum. Breeding cages were fed a high fat breeder chow. Mice were maintained and all procedures were performed within the Columbia University Institute of Comparative Medicine, which is fully accredited by the Association for Assessment and Accreditation of Laboratory Animal Care. All protocols were approved by the Columbia Institutional Animal Care and Use Committee.

### Genotyping

DNA was extracted from tail or ear clippings using the Kapa Mouse Genotyping Standard kit (KAPA Biosystems) and stored at -20°C. PCR was performed with 2x MyTaq HS Mix (Bioline), using the following Hnrnpu primers: FWD= 5’-GTCCGTTCTGCAGCAGCACT-3’, REV= 5’-TTACCTCCCGCCTGCTGTTG-3’. This amplifies a 745-bp product from the WT allele and a 632-bp product from the mutant allele.

### Primary neuronal culture

P0 pups were tail sampled, weighed and genotyped for the *Hnrnpu* 113-bp deletion. Mutant and wildtype pups were decapitated and cortex and hippocampus were separately dissected in cold Hibernate A (Thermo Fisher). Tissue was diced into smaller pieces and dissociated in a solution containing pre-warmed Hibernate A, papain and DNase for 20 min at 37°C. Dissociated tissue was then centrifuged at 300 g for 5 min at room temperature (RT), resuspended in pre-warmed Hibernate A, and triturated to further dissociate. Undissociated tissue was allowed to settle to the bottom of the tube, and the single cell suspension was transferred to a new tube and centrifuged at 300 g for 5 min at RT. The cell pellet was resuspended in complete medium containing Neurobasal A (Thermo Fisher), B27 Plus (Thermo Fisher), 1% FBS, Hepes, Glutamax and Penn/Strep. Cell viability and counts were obtained using a trypan blue exclusion assay, then further resuspended to the desired cell concentration using complete medium supplemented with laminin (5 ug/ml). Both cortical and hippocampal cells were plated on PDL-coated 12 mm coverslips in a 24-well dish at a density of 200,000 cells. Complete medium was changed the following morning to Neurobasal A, B27 Plus, Hepes, Glutamax and Penn/Strep, and 50% medium changes were subsequently performed every other day.

### Human induced pluripotent stem cell and cortical organoid cultures

CRISPR-Cas9 editing was performed on the PGP1 hiPSC line (Synthego) targeting exon 2 of human *HNRNPU*. Two heterozygous mutant clones that result in a frame shift and subsequent premature stop codon were selected and confirmed by Sanger sequencing. HET1 (clone D11) contains a 1bp duplication and HET2 (clone M20) contains a 10bp deletion. Karyotyping (Karyostat) and pluripotency analysis (PluriTest) were completed. Upon receipt, cells were expanded and banked using mTESR+ and geltrex coated plates.

Cortical organoids were generated using previously described Park protocol with limited alterations (https://www.ncbi.nlm.nih.gov/pmc/articles/PMC6402040/). Briefly, 9,000 live cells were plated on day 0 into ultra low adherent 96 well plates in neural induction (‘NI’) media (DMEM-F12 with KSR (15%), MEM-NEAA (1%), Glutamax (1%), B-Mercaptoethanol (100 uM), LDN-193189 (100 nM), SB431452 (10 uM), XAV939 (2 uM) supplemented with 5% FBS and 50 uM Rock-inhibitor. On day 2, media was removed and replaced with NI media with 50 uM Rock-inhibitor. Then, media was replaced without Rock-inhibitor every other day through day 10. On day 10, six to ten embryoid bodies were placed per well into ultra-low adherent 6-well plates and media was changed to Neural Differentiaton (‘ND’) media (DMEM-F12/Neurobasal with MEM-NEAA (.5%), Glutamax (1%), B-Mercaptoethanol (50 uM), Insulin (.125%), N2 (0.5%) and B27 without Vitamin A (1%). Plates were thereafter incubated on an orbital shaker set to 80RPM at 37C and media was changed every two days through day 18. On day 18, media was replaced with ND media with B27 including Vitamin A and additional neurotrophic factors (BDNF (20 ng/ul), cAMP (125 ug/ul), Ascorbic Acid (200 uM). Media was replaced every 4-5 days for the remainder of the experiments.

### Immunocytochemistry

On day in vitro 9 (DIV9), mouse primary cortical and hippocampal cells were washed 2x with 1X PBS, fixed for 15 min in 4% paraformaldehyde at RT, and again washed 2x with 1X PBS. Cells were incubated in a staining solution comprised of 5% donkey serum, 1% BSA, 0.3% TritonX-100 in 1X PBS for 15 min at RT, then subsequently incubated in the primary antibody diluted in the staining solution for 2 hr. at RT. Cells were washed 4x with 1X PBS, 0.2% TritonX-100, incubated with the fluorophore conjugated secondary antibody in staining solution for 30 min at RT then washed 4x with 1X PBS, 0.2% TritonX-100. Coverslips were mounted using ProLong-Antifade with DAPI on Superfrost Plus Microscope slides and allowed to dry in the dark prior to imaging. Imaging was performed using the Zeiss Axio Observer.Z1 Fluorescence Motorized Microscope and associated Zen2 Pro imaging software. Downstream image processing was performed using Adobe Photoshop, using auto-brightness and contrast for each individual channel and merged image. Primary antibodies used include: Mouse anti-Map2 at 1:500 (Sigma M4403), mouse anti-GFAP at 1:100 (Abcam ab10062), mouse anti-Gad67 at 1:500 (Millipore MAB5406), mouse anti-Satb2 at 1:100 (Abcam ab51502), rat anti-Ctip2 at 1:250 (Abcam ab18465), rabbit anti-HNRNPU at 1:500 (Abcam 20666). Secondary antibodies include: 488 and 568 Alexa Fluorophore conjugated donkey anti-mouse, donkey anti-rabbit and donkey anti-rat (Invitrogen), at 1:1000 dilution.

### Western blotting

Dissected tissue was snap frozen in liquid nitrogen and stored at -80°C until time of extraction. Tissue was thawed on ice and homogenized using a motorized pestle in RIPA buffer containing both protease and phosphatase inhibitor cocktails (Roche). Lysis was completed for 15 minutes on ice. Samples were subsequently centrifuged at full speed for 20 minutes at 4°C. The resulting supernatant was collected and protein was quantified using the BCA method (Pierce) with BSA as a standard. All western blots were performed using the Novex NuPAGE system (Invitrogen). Protein lysates were diluted in LDS sample buffer and reducing agent, and heated at 70°C for 10 min. Using the Xcell SureLock Mini Cell gel box, a total of 5 ug of reduced protein lysates were loaded onto a 4-12% gradient Bis-Tris gel in 1X SDS Running buffer (Invitrogen) supplemented with the NuPAGE antioxidant, and ran at 180 V for 1-1.5 hrs. Using the Xcell II Blot Module, proteins were subsequently transferred to a 0.2 um methanol-activated PVDF membrane at 30 V for 1.25 hrs at 4°C in Transfer buffer containing 20% methanol. Membranes were blocked for 1 hr at RT in 5% milk, then incubated overnight in the hnRNP U primary antibody at 1:1000 (Rabbit polyclonal against C-terminus: Abcam ab20666; Rabbit monoclonal against N-terminus: Abcam ab180952) diluted in 5% BSA. Blots were washed 3x for 10 min in PBST, incubated at RT for 1 hr in a secondary HRP-conjugated anti-rabbit (at 1:10,000) diluted in 5% BSA, then further washed 3x for 10 min in PBST. Proteins were incubated for 5 min in a standard ECL substrate (Pierce) and developed with either a Kodak X-OMAT 2000A Processor or iBright FL1000 Imaging system (Invitrogen). For a loading control, blots were subsequently incubated in an HRP-conjugated b-Actin secondary at 1:1000 (Santa Cruz #sc-47778) diluted in 5% BSA for 1 hr at RT, then washed and developed as previously described. Densitometry analysis was performed using the iBright FL1000 imager. Specifically, the Local Background Corrected Density, LBCD, (background-corrected volume/area) of each hnRNP U-probed sample was first normalized to the corresponding LBCD of β-Actin to generate an hnRNP U/β-Actin ratio. WT and HET ratios were further divided by the average WT hnRNP U/β-Actin ratio and plotted individually.

### qRT-PCR

Human iPSCs and cortical organoids: RNA was extracted from stem cells (1 confluent well of 6 well plate per sample), day 10 embryoid bodies (EBs) and D45-47 cortical organoids (3 pooled EBs or COs per sample) using RNeasy Mini kit (Qiagen). Approximately 0.5-1ug of RNA was used as input to generate cDNA using Superscript IV Reverse Transcriptase (ThermoFisher) with random hexamer priming. The following TaqMan probes (ThermoFisher) were used: *ATP5F1* (Hs01076982_g1, FAM), *HNRNPU* (Hs00244919_m1, FAM) and primer limited *GAPDH* (Hs99999905_m1, VIC). *GAPDH* was ran in duplex with *ATP5F1* and *HNRNPU*. TaqMan Fast Universal Master Mix 2X (Applied Biosystems) was performed in a total volume of 10uL per well with 0.5uL of each probe and approximately 10ng of template cDNA. TaqMan assays were run on a QuantStudio 5 Real-Time PCR System (Applied Biosystems) using the comparative Ct method.

Mouse cerebral cortex: Cortical tissue was collected and immediately stored in RNALater Stabilization Solution (Qiagen) at 4°C. After 24 hrs., the RNALater was subsequently removed, and samples were stored long term at -80°C. For RNA extraction, tissue was first mechanically homogenized using a motorized pestle in RLT buffer (Qiagen) supplemented with b-mercaptoethanol, then further homogenized using a QIAshredder spin column (Qiagen). RNA was extracted using the RNeasy Mini kit (Qiagen) per protocol instructions, and the resulting RNA concentration and purity were assessed using a NanoDrop. A total of 2.5 ug of RNA was used for the reverse transcription reaction, which was performed using the SuperScript IV First-Strand Synthesis System (ThermoFisher) with random hexamer priming. Resulting cDNA was used as input into pre-validated TaqMan Gene Expression Assays (ThermoFisher) and run with the TaqMan Fast Universal PCR MasterMix 2x (Applied Biosystems). The following TaqMan probes were purchased from ThermoFisher: mHnrnpu Mm00469329_m1 (spans exons 1-2) and mCyc1 Mm00470540_m1 (spans exons 1-2). A total of six biological replicates were evaluated. TaqMan assays were run on a QuantStudio 5 Real-Time PCR System (Applied Biosystems) using the comparative Ct method.

All qRT-PCR analyses were performed using QuantStudio Design and Analysis Software v1.2 and Microsoft Excel. To analyze for gene expression differences, raw Ct values were first averaged across technical replicates, which ranged from 2-4 for each sample. For samples with a Ct standard deviation >0.3, a technical replicate was filtered out if its Ct value was one standard deviation above or below that of the mean of the replicates. We required samples to have at least technical duplicates to be considered in the final analysis. Delta Ct values were determined by calculating the difference between mean Ct values for experimental samples and corresponding loading controls (*Cyc1* for mouse cortex; *GAPDH* and *ATP5F1* for hiPSCs and hCOs). Delta delta Ct values for both HET and WT samples were then calculated using the average WT delta Ct from the corresponding time point and experimental batch, and then were independently plotted along with SEM. Statistical analyses were performed using Welch’s two-sample t-test.

### Morphological studies

Brains were extracted from P0 pups and fixed in Bouin’s solution overnight at room temperature. Fixed brains were embedded in paraffin with service provided by Columbia University’s Molecular Pathology Core Facility. Coronal sections in 5 μm thickness were obtained using a microtome (Leica RM2125RT) and subjected to hematoxylin and eosin (H&E) staining. Briefly, the slices were deparaffinized in xylene and rehydrated with ethanol and water. Slices were then stained in hematoxylin, counterstained with eosin and subsequently dehydrated with ethanol and xylene. Stained slices were mounted with coverslips using Permount (Fisher Chemical). Images were acquired using a Nikon Eclipse E800 Microscope packaged with NIS-Elements DV.4.51.00 imaging software. For presentation purpose, full size images were subjected to automatic brightness and contrast adjustment using Adobe Photoshop. The morphological analysis was performed by a blinded reviewer. Brain measurements were collected using ImageJ. The measurements were normalized to respective pup body weight. Student’s T-test was performed on each set of measurements and Bonferroni corrected for multiple comparisons.

### Pup developmental milestones

On postnatal day 2, pups were tattooed on the bottom of their paws for identification using a 25-30 G needle and Ketchum tattoo ink, following the AIMS tattoo chart. The following developmental milestone studies were completed on P4, P6, P8, P10 and P12 within a 3 min time window for each test subject.

Righting reflex: the latency to flip over from supine to prone position on all fours. Pups were gently placed on their backs on a hard surface and released. A stopwatch was used to measure the total time for each pup to right itself. The cutoff latency was 30 s.

Negative geotaxis: the latency to face upwards from a downward-facing start position on an inclined mesh screen. Pups were placed at a downward facing position on a 45° inclined screen and released. A stopwatch was used to record the time it takes for each pup to turn 90° then 180°. The cutoff latency was 30 s. If the pup failed the trial by falling while turning, they were scored the maximum cutoff latency time of 30 s.

Vertical screen: the latency to fall from a vertically-positioned wire mesh screen. Pups were positioned to grasp the screen. Using a stopwatch, the time until fall was recorded. The minimum and maximum latency allowed was 1 s and 30 s, respectively.

For each developmental test, every pup underwent two successive trials that were subsequently averaged and used in the downstream analyses. Throughout the testing period, the experimenter was blinded to genotype. The mean and standard error of the mean (SEM) for both genotypes were plotted across all tested timepoints. Permuted MWU p-values were reported for each time point. Briefly, WT and HET data were pooled, randomly sampled and MWU p-values calculated 10,000 times. Permuted MWU p-values were calculated as the total number of randomly-sampled MWU p-values that fell below the actual MWU p-value divided by the total number of permutations (10,000).

### Pup ultrasonic vocalizations (USVs)

USVs were assessed on P3, P5, P7, P9 and P11. Each pup was gently removed from the nest and placed in a small, plastic container containing a 0.5 cm layer of fresh bedding. The cage lid was immediately returned to avoid irritating the dam and remaining pups in the nest. The container holding the pup was placed immediately into a sound-attenuating environmental chamber (Med Associates, St. Albans, VT, USA). After a 3 min recording, each pup was marked and returned to the nest. Ultrasonic vocalizations were recorded with an Ultrasound Microphone (Avisoft UltraSoundGate condenser microphone capsule CM16, Avisoft Bioacoustics, Berlin, Germany) sensitive to frequencies of 10-180 kHz and using the Avisoft Recorder (Version 4.2) software. Sampling rate was 250 kHz, format 16 bit. Ultrasonic vocalizations were analyzed using Avisoft SASLab Pro software (Avisoft Bioacoustics). Spectrograms were generated for each 1 min audio file, with an FFT-length of 512 points and a time window overlap of 75% (100% Frame, Hamming window). The spectrogram was generated at a frequency resolution of 488 Hz and a time resolution of 1 ms. A lower cut-off frequency of 15 kHz was used to reduce background noise outside the relevant frequency band to 0 dB. Calls were inspected visually and manually labelled. Summary statistics were generated by Avisoft SASLab Pro and analyzed using Prism. All calls emitted over the 3 min recordings were quantified. For the qualitative analysis, one 1 min file (out of three 1 min files) that included the most USVs were analyzed for each mouse. Mean and SEM were plotted, and permuted MWU p-values were reported for each time point as described for pup developmental milestone tests.

### Electroencephalography (EEG)

Video EEGs were performed on 6-to 8-week-old adult mice. Mice were anesthetized with tribromoethanol (250 mg/kg delivered via intraperitoneal injection, Sigma Aldrich cat# T48402), and three small burr holes were drilled through the skull 2 mm lateral to the midline (1mm rostral to the bregma on both sides and 2mm caudal to the bregma on the left). One hole was also drilled over the cerebellum as a reference. Four Teflon-coated silver wires soldered onto pins of a microconnector (Mouser electronics cat# 575-501101) were placed in between the dura and brain. A dental cap was applied on top. Each mouse was provided the post-operative analgesic Carprofen (5 mg/kg subcutaneous Rimadyl) and allowed a recovery period of at least 48 hours prior to recording. Signal was obtained on either a Grael II EEG amplifier (Compumedics) or Natus Quantum amplifier (Natus Neuro). Data was analyzed with either Profusion 5 (Compumedics) or NeuroWorks (Natus Neuro). Differential amplification recordings were recorded pairwise between all three electrodes and the reference, resulting in 6 total channels for each subject. Mouse behavior was captured throughout the recording period through video using a Sony IPELA EP550 camera with infrared light for dark recordings. Each mouse was recorded for 24-48 hrs. continuously.

### Electroconvulsive threshold (ECT) studies

All tests were performed on 6-to 8-week-old mice. Transcorneal electrodes were used to deliver a predefined stimulus with the Ugo Basile Model 7801 electroconvulsive device. High frequency (HF) electroshock was performed with the following fixed settings: 1.6 ms pulse width, 0.2 s shock duration and 299 Hz pulse frequency with variable settings of 4-12 mA amplitude. The individual threshold for each mouse was determined by testing in 0.5 mA intervals on sequential days until the threshold was reached. The behavioral endpoint evaluated was a maximal tonic hindlimb extension seizure, which often start with tonic extension of the forelimbs that evolves into full tonic hindlimb extension. The overall stimulus is calculated as the iRMS (integrated root mean square, or the integrated area under the curve) using the following equation: sq. root frequency (Hz) x pulse width (ms) x duration (s) x amplitude (mA).

### Adult behavioral tests

Elevated Plus Maze test (EPM): This classic test for anxiety-like behavior is based on rodents’ innate fear for height and open space. The elevated plus-maze consists of two open arms (30 cm x 5 cm) and two closed arms (30 × 5 × 15 cm) extending from a central area (5 × 5 cm). Photo beams embedded at arm entrances register movements. Room illumination was approximately 5 lux. The test begins by placing the subject mouse in the center, facing a closed arm. The mouse is allowed to freely explore the maze for 5 min. Time spent in the open arms and closed arms, the junction, and number of entries into the open arms and closed arms, are automatically scored by the MED-PC V 64bit Software (Med Associates). At the end of the test, the mouse is gently removed from the maze and returned to its home cage. The maze is cleaned with 70% ethanol and wiped dry between subjects.

Open Field exploratory activity: The open field test is the most commonly used general test for locomotor activity. Each mouse is gently placed in the center of a clear Plexiglass arena (27.31 × 27.31 × 20.32 cm, Med Associates ENV-510) lit with dim light (∼5 lux), and is allowed to ambulate freely for 60 min. Infrared (IR) beams embedded along the X, Y, Z axes of the arena automatically track distance moved, horizontal movement, vertical movement, stereotypies, and time spent in center zone. At the end of the test, the mouse is returned to the home cage and the arena is cleaned with 70% ethanol followed by water, and wiped dry.

Catwalk: Free-pace walking was evaluated using the Catwalk XT system (Noldus Information Technology) which consists of an illuminated walled glass walkway (130 cm x 10 cm) and a high-speed camera underneath. Light is reflected and illuminates the stimulus (footprint) when downward pressure is applied. Walking patterns are captured with a high-speed camera mounted underneath the walkway. The experiment was done with dim room illumination (30 lux). The mouse is allowed to traverse the walkway as many times as needed to obtain at least 3 compliant runs (runs with a speed variation under 80% in 20 seconds or less). Pilot experiments using a 60% speed variation limit (most common in the literature) proved to be too stringent for most heterozygous mice. Parameters automatically collected by the software include, but are not limited to, paw statistics, intensity measures, stride length, width, base of support, distance between ipsilateral prints, cadence, % limb support, regularity index, speed, and speed variation. A highly trained experimenter visually inspected all automatically scored runs, and manually classified any prints that were too ambiguous for the software to identify accurately. The walkway is cleaned with paper towel moistened with 70% ethanol and wiped dry between trials.

Acoustic startle response: Acoustic startle response was tested using the SR-Laboratory System (San Diego Instruments, San Diego, CA). Test sessions began by placing the mouse in the Plexiglass holding cylinder for a 5-min acclimation period. For the next 8 min, mice were presented with each of six trial types across six discrete blocks of trials, for a total of 36 trials. The inter-trial interval was 10–20 s. One trial type measured the response to no stimulus (baseline movement). The other five trial types measured startle responses to 40 ms sound bursts of 80, 90, 100, 110 or 120 dB. The six trial types were presented in pseudorandom order such that each trial type was presented once within a block of six trials. Startle amplitude was measured every 1 ms over a 65 ms period beginning at the onset of the startle stimulus. The maximum startle amplitude over this sampling period was taken as the dependent variable. Background noise level of 70 dB was maintained over the duration of the test session.

Fear Conditioning: This is a classic test for conditioned learning. Training and conditioning tests are conducted in two identical chambers (Med Associates, E. Fairfield, VT) that were calibrated to deliver identical foot shocks. Each chamber was 30 cm × 24 cm × 21 cm with a clear polycarbonate front wall, two stainless side walls, and a white opaque back wall. The bottom of the chamber consisted of a removable grid floor with a waste pan underneath. When placed in the chamber, the grid floor connected with a circuit board for delivery of scrambled electric shock. Each conditioning chamber is placed inside a sound-attenuating environmental chamber (Med Associates). A camera mounted on the front door of the environmental chamber recorded test sessions which were later scored automatically, using the VideoFreeze software (Med Associates, E. Fairfield, VT). For the training session, each chamber is illuminated with a white house light. An olfactory cue is added by dabbing a drop of imitation almond flavoring solution (1:100 dilution in water) on the metal tray beneath the grid floor. The mouse is placed in the test chamber and allowed to explore freely for 2 min. A pure tone (5 kHz, 90 dB) which serves as the conditioned stimulus (CS) is played for 30 s. During the last 2 s of the tone, a foot shock (0.5 mA) is delivered as the unconditioned stimulus (US). Each mouse received three CS-US pairings, separated by 90 s intervals. After the last CS-US pairing, the mouse is left in the chamber for another 120 s, during which freezing behavior is scored by the Video Freeze software. The mouse is then returned to its home cage. Contextual conditioning is tested 24 h later in the same chamber, with the same illumination and olfactory cue present but without foot shock. Each mouse is placed in the chamber for 5 min, in the absence of CS and US, during which freezing is scored. The mouse is then returned to its home cage. Cued conditioning is conducted 48 h after training. Contextual cues are altered by covering the grid floor with a smooth white plastic sheet, inserting a piece of black plastic sheet bent to form a vaulted ceiling, using near infrared light instead of white light, and dabbing vanilla instead of banana odor on the floor. The session consisted of a 3 min free exploration period followed by 3 min of the identical CS tone (5 kHz, 90 dB). Freezing is scored during both 3 min segments. The mouse was then returned to its home cage. The chamber is thoroughly cleaned of odors between sessions, using 70% ethanol and water.

Y maze: The Y-maze is a standard test for assessing short term memory in mice, based on the mouse’s natural tendency to explore novel locations. Memory impairment is indicated by failing to spend more time exploring the novel arm than the familiar arms. The test is conducted in the Y maze (Maze Engineer) consisting of three arms of equal arm lengths (35 cm), arm lane width (5 cm), wall height (10 cm). One arm is the start arm, with a “=” sticker velcroed on the wall, to the end of the arm. The two stickers (bus and plane) are velcroed on the wall at the end of the other two arms. The placement of the stickers was counterbalanced across animals. The novel arm preference test consists of two trials. In trial 1, each mouse is placed in the designated start arm and allowed access the start arm and the one other arm for 10 min. The third arm is blocked with an opaque door. At the conclusion of trial 1, the mouse was placed in a temporary holding cage for 10 min. For trial 2, the subject mouse was returned to the start location, and allowed to explore all arms for 5 min. A camera mounted above the maze and interfaced with the Ethovision software (Noldus Information Technology) automatically records distance traveled, arm entries, and time spent in each arm. The maze was cleaned with 50% ethanol and allowed to dry between trials and between animals.

Morris water maze: Spatial learning and reversal learning were assessed in the Morris water maze using procedures and equipment as previously described^71^. The apparatus was a circular pool (120 cm diameter) filled 45 cm deep with tap water rendered opaque with the addition of non-toxic white paint (Crayola, Easton, PA). Distal room cues were door, chairs, computers, and proximal cues are two 20cm x 20cm stickers. Trials were recorded and automatically scored by Ethovision 12 (Noldus Information Technology). Acquisition training consisted of 4 trials a day for 5 days. Each training trial began by lowering the mouse into the water close to the pool edge, in a quadrant that was either right of, left of, or opposite to, the target quadrant containing the platform. The start location for each trial was alternated in a semi-random order for each mouse. The hidden platform remained in the same quadrant for all trials during acquisition training for a given mouse, but varied across subject mice. Mice were allowed a maximum of 60 s to reach the platform. A mouse that failed to reach the platform in 60 s was guided to the platform by the experimenter. Mice were left on the platform for 15 s before being removed. After each trial, the subject was placed in a cage lined with absorbent paper towels and allowed to rest under an infrared heating lamp for 60 s. Two hours after the completion of training on day 5, the platform was removed and mice were tested in a 60 s probe trial. Parameters recorded during training days were latency to reach the platform, total distance traveled, and swim speed. Time spent in each quadrant and number of crossings over the trained platform location and over analogous locations in the other quadrants were used to analyze probe trial performance.

### UV crosslinking immunoprecipitation and sequencing (CLIP-seq) analysis

CLIP-seq experiments were conducted with hearts from two-week-old WT mice using anti-hnRNP U (A300-690A, Bethyl Laboratories). Sample preparation, crosslinked-RNA recovery, library preparation and sequencing were performed according to published protocols (Moore et al 2014). After linker sequence trimming and duplication collapsing, CLIP reads were aligned to mouse genome (mm10) by Novoalign, and unique tags were clustered. Distribution of CLIP tags to different genomic regions were determined. We obtained 822,984 unique tags for hnRNP U with a majority (63%) of these tags mapped to introns, 8% mapped to exons and the remaining tags mapped to promoter, 5’ and 3’ UTRs and intergenic regions.

### Single-cell RNA-sequencing and data integration

Neocortical and hippocampal tissue was dissected from postnatal day 0 pups and subjected to a papain dissociation. Following papain dissociation and tissue trituration, all neocortical and hippocampal samples were filtered through a 40 μm cell strainer to enrich for single cells in the resulting suspension. Cell viability was subsequently assessed, with a cutoff of 70% or greater to be used for sequencing. Single cell RNA-seq libraries were constructed using the 10X Chromium Single Cell 3′ Reagent Kits v2 according to manufacturer descriptions, and samples were sequenced on a NovaSeq 6000. Reads were aligned to the mm10 genome using the 10X CellRanger pipeline with default parameters to generate the feature-barcode matrix.

We used Seurat v3 to perform downstream QC and analyses on feature-barcode matrices^37,38^. We removed all genes that were not detected in at least 4 cells. We further removed cells with fewer than 1,000 genes or more than 5,000 genes detected. For cortical cells, we removed all cells with greater than 8% of reads mapping to mitochondrial genes. For hippocampal cells, we removed all cells with greater than 15% of reads mapping to mitochondrial genes. The filtered matrices were log-normalized and scaled to 10,000 transcripts per cell. We used the variance-stabilizing transformation implemented in the FindVariableFeatures function in order to identify the top 2,000 most variable genes per sample. We used Seurat’s data integration method to harmonize gene expression across datasets prior to clustering. We first identified anchors between samples in each dataset using the FindIntegrationAnchors function, which uses canonical correlation analysis (CCA) to identify pairwise cell correspondences between samples. We then computed an integrated expression matrix using these anchors as input to the IntegrateData function.

Next, we used linear regression to regress out the number of UMIs per cell and percentage of mitochondrial reads using the ScaleData function on the integrated expression matrices. We then performed dimensionality reduction using PCA. For each dataset, we selected the top 30 dimensions to compute a cellular distance matrix, which was used to generate a K-nearest neighbor graph. The KNN was used as input to the Louvain Clustering algorithm implemented in the FindClusters function. For clustering via Louvain, we chose a resolution parameter of 0.8. We visualized the cells using UMAP via the RunUMAP function.

To annotate and merge clusters, we performed differential gene expression analysis on the integrated expression values between each cluster using the default parameters in the FindMarkers function, which implements a Wilcoxon test and corrects p-values via Bonferroni correction. Additionally, we visualized the expression of canonical marker genes aggregated from previous single-cell publications^72–75^. Clusters representing microglia and smooth muscle cells were excluded given limited (5-10) cells represented from these cell types.

### Differential gene expression analysis

We performed cell-type-specific differential gene expression analysis using MAST^76^, as implemented in Seurat’s FindMarkers function, in order to identify genes dysregulated between mutant and wildtype cells. We excluded all non-coding genes, genes encoding ribosomal proteins, and pseudogenes from our analysis to reduce the multiple testing burden. For each cell type, we fit a linear mixed model that included the gene detection rate (ngeneson) and gender as latent variables: zlm(∼ngeneson + gender)

We corrected the p-values using the Benjamini-Hochberg FDR method. We considered genes with a log2(fold change) value of at least 0.14 (10% difference) and FDR < 0.05 as differentially expressed. We performed gene ontology analysis using g:Profiler(Raudvere et al., 2019), using all tested genes per cell type as a background set. P-values were generated using Fisher’s Exact Test and corrected via FDR.

For the burden analysis, we down sampled the data to include 300 cells per population. To increase power, we pooled cells from male and female samples. We ran the differential expression analysis as described above, removing gender as a latent variable.

For disease gene enrichment analyses, human homologs for all tested mouse genes were obtained using biomaRt. Significant and nonsignificant mouse genes were annotated based on the respective human homolog disease gene status. Genes without human homologs were not used in the analysis. The epilepsy-associated gene list was based on a prior publication^77^. Autism genes were based on SFARI genes with gene scores 1, 2, 3 and S on SFARI.org^78^. Confirmed monoallelic developmental delay genes (obtained in 2019) were obtained from the Deciphering Developmental Disorders study^79^. A Fisher’s exact test was performed on significant downregulated and upregulated DEGs compared to all non-significant DEGs.

### Transcriptomic reversal

We queried the Connectivity Map (clue.io) to identify compounds most likely to reverse disease expression signatures^44,45^. Our disease expression signatures included the top 150 downregulated genes per query. We compared the compounds predicted to reverse the expression profile derived from excitatory cells in the subiculum as well as the expression profile that would have been recovered via bulk RNA-sequencing. To generate this second signature (i.e. the pseudo-bulk signature), we performed differential gene expression between mutant and wild type cells using MAST without factoring in cell types. We considered compounds that achieved a Connectivity Score of less than -90 as most likely to reverse the disease signatures.

## Acknowledgements

The authors thank Megha Sah, Virginia Osasumwen and Daniel Krizay for thoughtful discussions and expertise on various experiments, along with Sahar Gelfman for his input on data analysis. We also thank Erin Bush and the Columbia Sulzberger Genome Center for their assistance in performing single-cell sequencing, and Andrew Butler for helpful discussions regarding single-cell RNA-sequencing clustering and differential gene expression analysis. The authors are also grateful to the Robbins Family for their continued advocacy and financial support of this work. This work was further supported by: NIH grant R37 NS031348, T32 Training Grant TL1TR001875.

## Author contributions

Conceptualization: S.A.D., R.S.D., M.J.B., W.N.F., D.B.G. Methodology: S.A.D., R.S.D., S.C., M.Y., M.J.B., W.N.F. Data acquisition: S.A.D., G.D.A.S., A.K.R., E.R., S.P., V.L., J.Y. Software: R.S.D. Formal Analysis: S.A.D., R.S.D., J.T., J.Y., M.Y., W.N.F. Interpretation: S.A.D., R.S.D., A.K.R., J.T., M.Y., M.J.B., W.N.F., D.B.G. Writing—Original Draft: S.A.D., R.S.D. Writing—Review & Editing: S.A.D., R.S.D., J.T., J.Y., S.C., M.Y., M.J.B., W.N.F., D.B.G. Visualization: S.A.D., R.S.D. Supervision: M.Y., M.J.B., W.N.F., D.B.G. Funding Acquisition: W.N.F., D.B.G

## Competing interests

D.B.G. is a founder of and holds equity in Praxis, serves as a consultant to AstraZeneca, and has received research support from Janssen, Gilead, Biogen, AstraZeneca and UCB. R.S.D serves as a consultant to AstraZeneca. S.C. serves a consultant for Q-State Biosciences, Inc

## Supplemental Figures

**Supplemental Figure 1.**
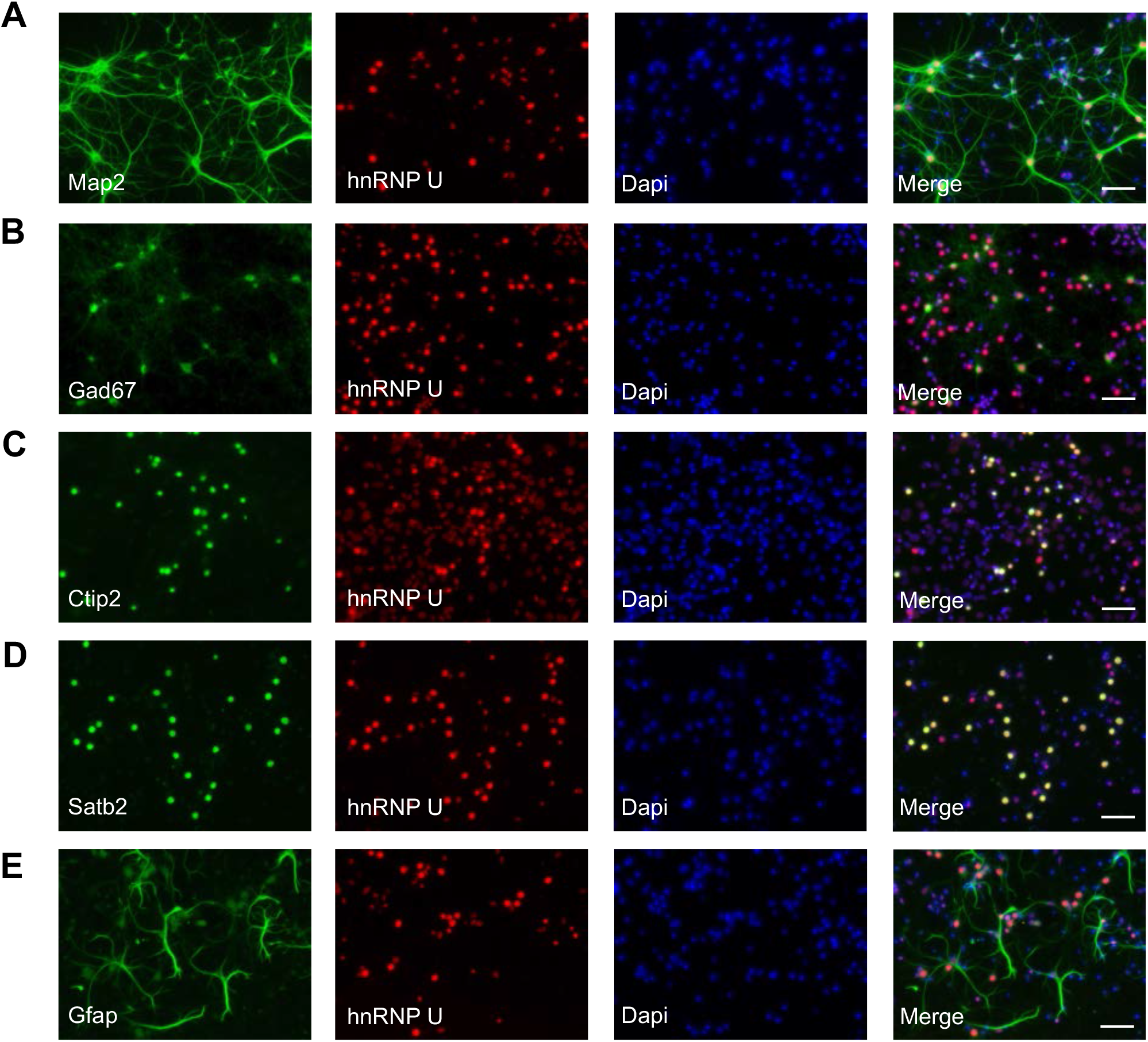
HnRNP U is expressed ubiquitously in neocortical cells. Primary neocortical cell cultures at day *in vitro* 9 showing hnRNP U (red) co-staining with nuclear marker Dapi (blue) and cell markers (green) **(A)** Map2 (neuronal), **(B)** Gad67 (inhibitory), **(C)** Ctip2 (deeper layer pyramidal neurons), **(D)** Satb2 (upper layer pyramidal neurons) and **(E)** Gfap (astrocytes). Scale bar= 50μm.

**Supplemental Figure 2.**
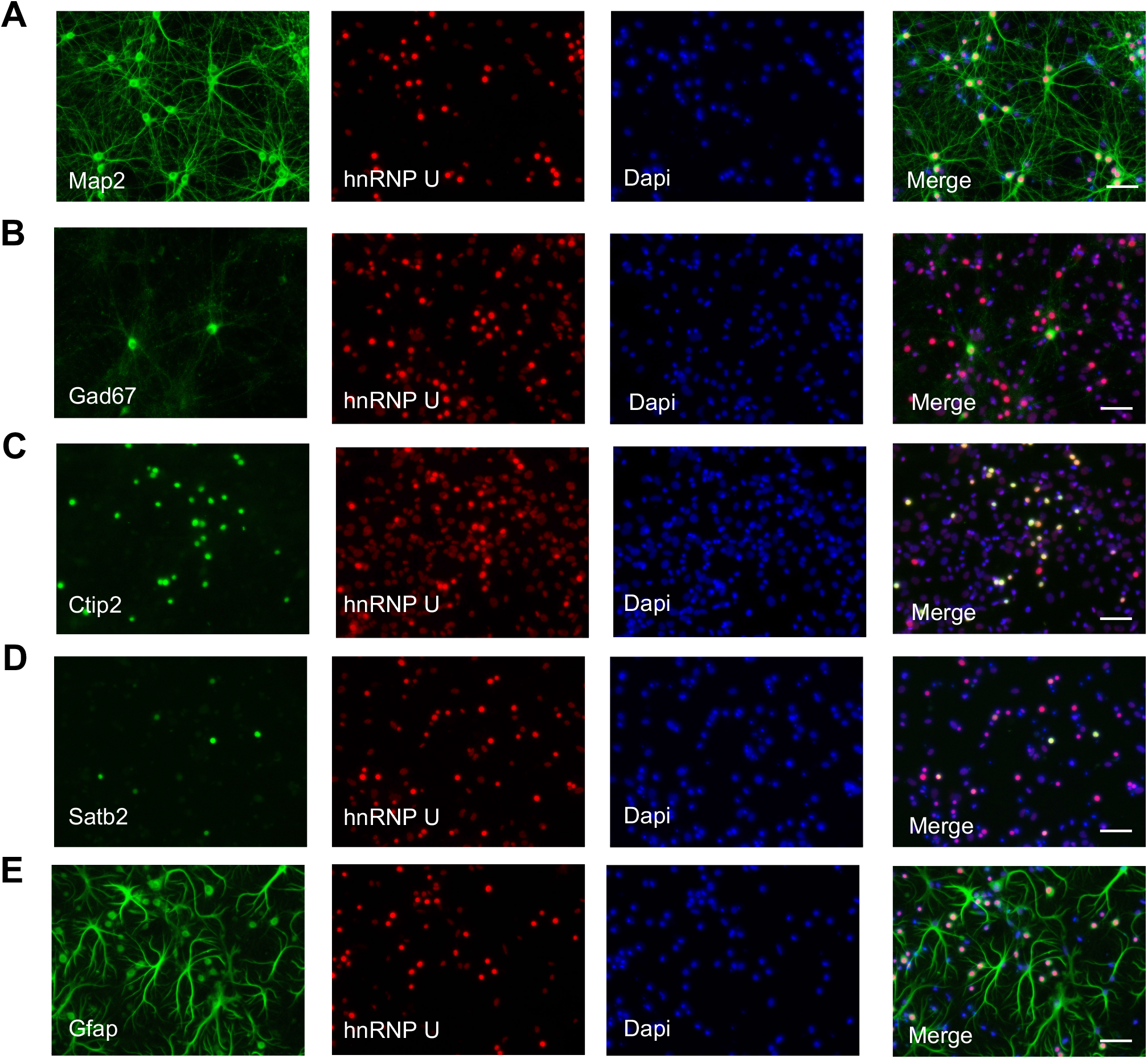
HnRNP U is expressed ubiquitously in hippocampal cells. hnRNP U (red) co-staining with nuclear marker Dapi and cell markers (green) **(A)** Map2 (neuronal), **(B)** Gad67 (inhibitory), **(C)** Ctip2 (deeper layer pyramidal neurons), **(D)** Satb2 (upper layer pyramidal neurons) and **(E)** Gfap (astrocytes). Scale bar= 50μm.

**Supplemental Figure 3.**
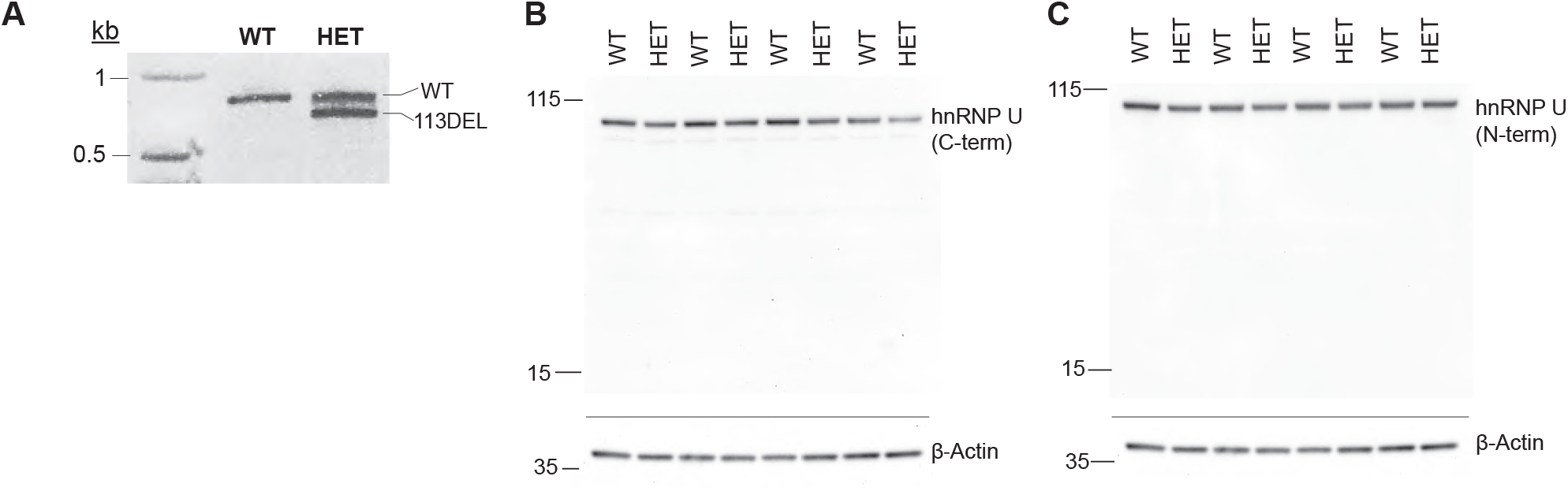
Generation and viability of a heterozygous *Hnrnpu* knockout mouse model. **(A)** A representative genotyping agarose gel showing the 113-bp deletion. **(B)** Western blot image using antibody targeting hnRNP U C-terminus **(C)** Western blot image using an antibody targeting hnRNP U N-terminus.

**Supplemental Figure 4.**
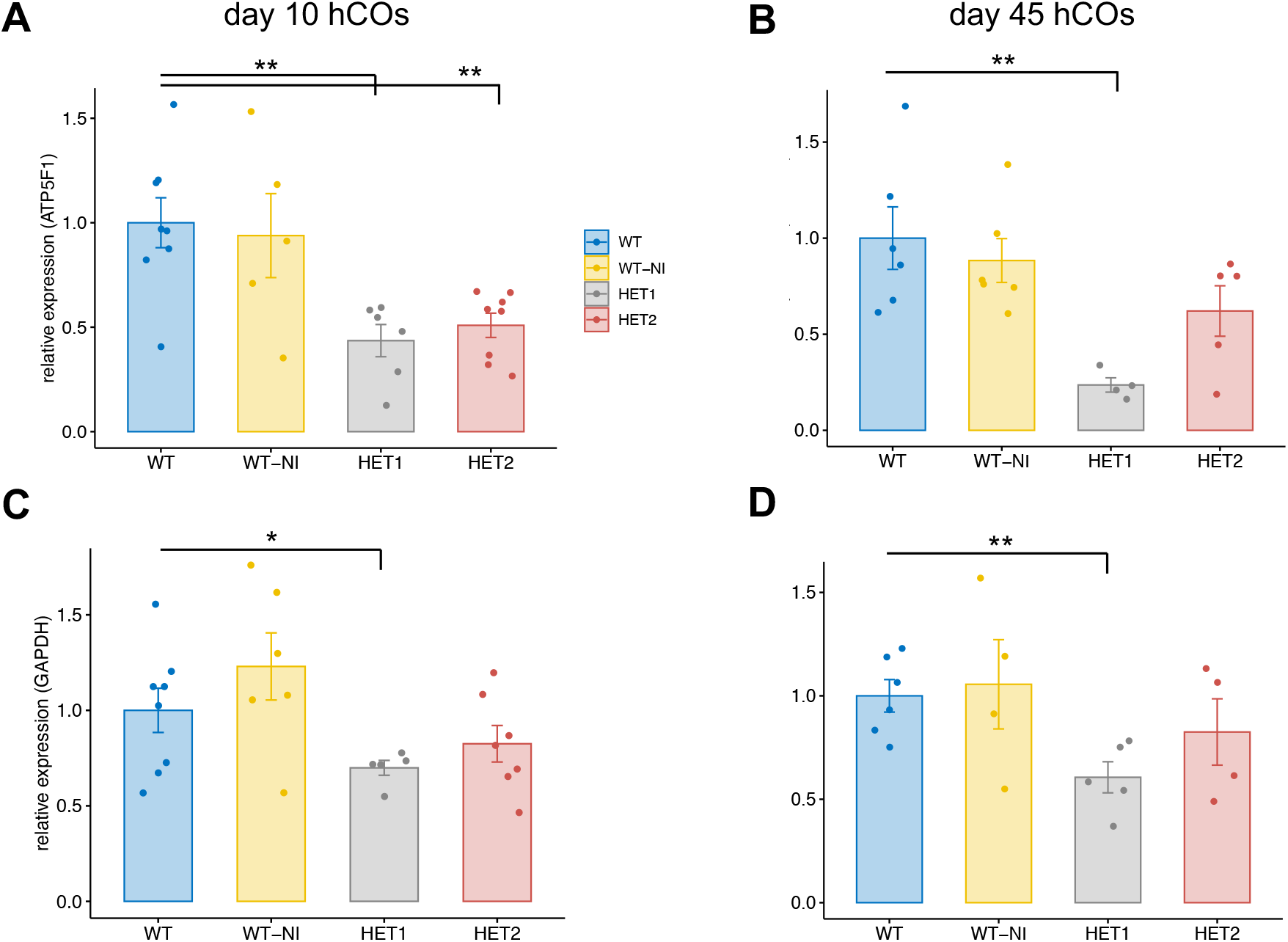
Variable *HNRNPU* expression in a heterozygous *HNRNPU* mutant human disease model. *HNRNPU* expression relative to the reference gene *ATP5F1* of **(A)** day 10 human cortical organoids (hCOs) (Welch’s t-test p-values: HET1=2.1×10^−3^, HET2=4.1×10^−3^) and **(B)** day 45 hCOs (Welch’s t-test p-values: HET1=4.7×10^−3^, HET2=0.10). *HNRNPU* expression relative to the reference gene *GAPDH* of **(C)** day 10 hCOs (Welch’s t-test p-values: HET1=0.04, HET2=0.27) and **(D)** day 45 hCOs (Welch’s t-test p-values: HET1=5.6×10^−3^, HET2=0.4). WT= WT isogenic control, WT-NI= WT non-isogenic control, HET1 = 1bp insertion in exon 2, HET2= 10bp deletion in exon 2. Statistics were only performed using the mutants and WT isogenic control. Error bars= SEM.

**Supplementary Figure 5.**
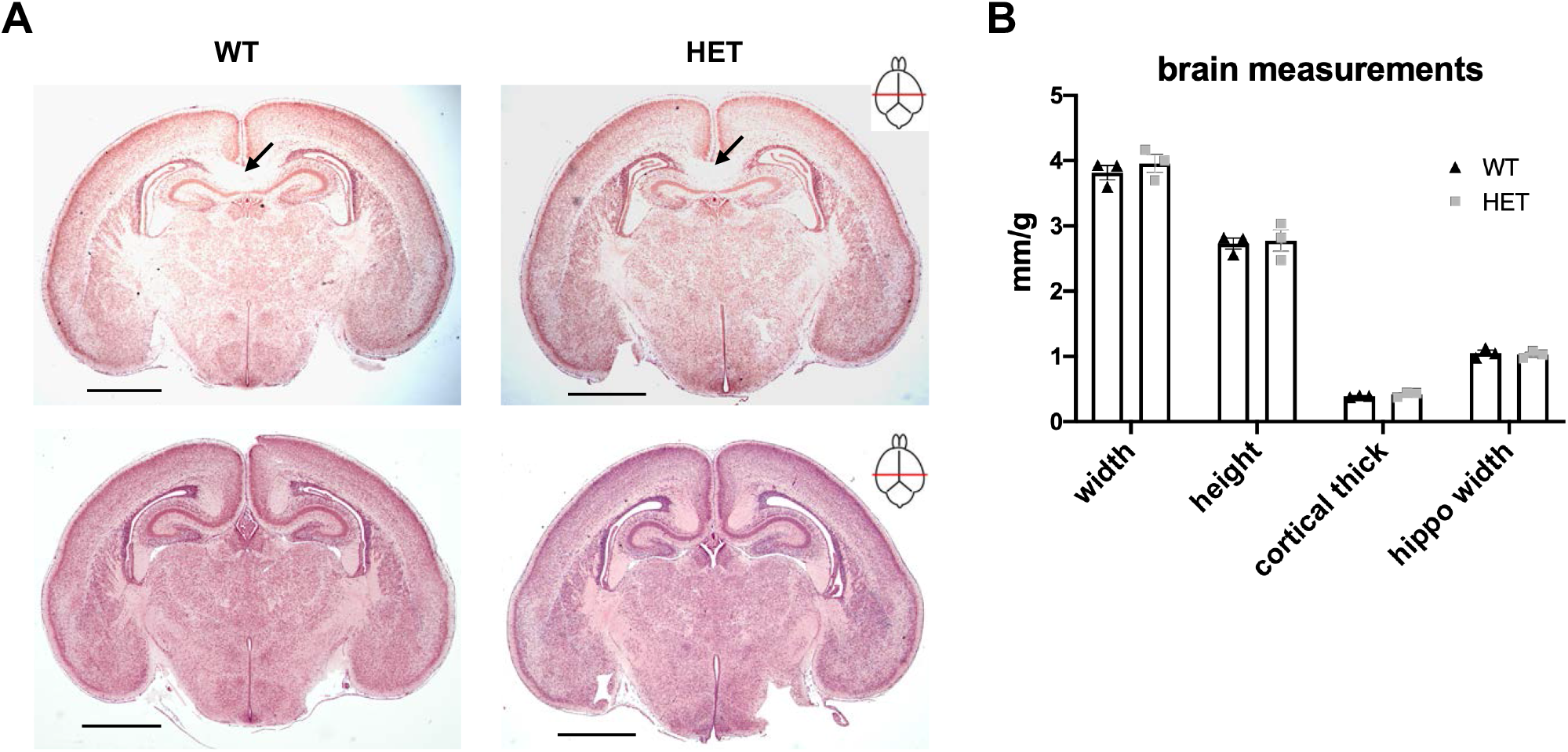
No overt morphological defects observed in *Hnrnpu*^+/113DEL^ mice. **(A)** Representative H&E-stained coronal sections of WT and HET brains at P0. Black arrows indicate the corpus callosum. Level of sections indicated by the respective cartoons. Scale bar= 1 mm. **(B)** Brain width, height, cortical thickness and hippocampal width were measured from caudal sections and normalized to respective body weight (n= 3 animals per genotype). Bonferroni-corrected T-test p> 0.99 for each test (width t= 0.79, height t= 0.25, cortical thickness t= 1.2, hippocampus width t= 0.39, df= 4). Error bars = SEM.

**Supplemental Figure 6.**
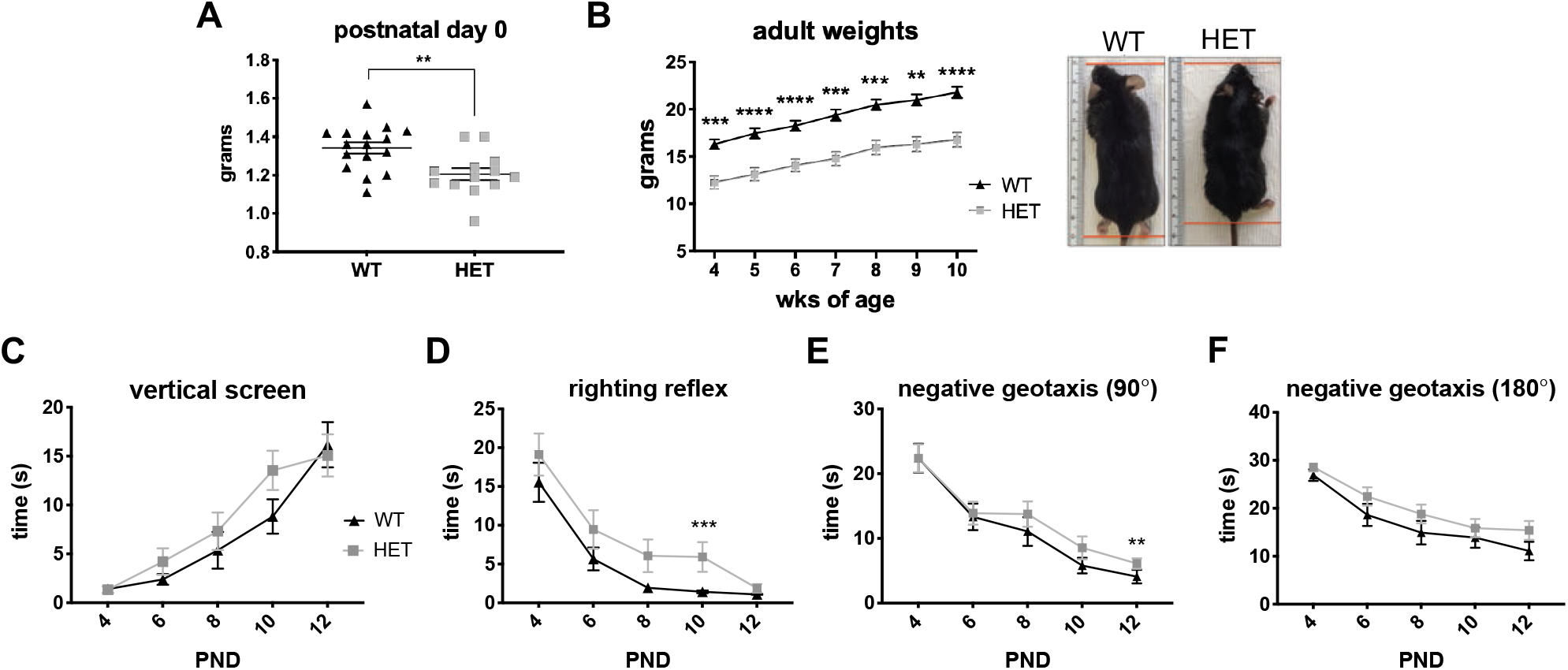
Delayed physical and sensorimotor development in *Hnrnpu*^+/113DEL^ mice. **(A)** Body weight of postnatal day 0 (P0) pups (n=16 WT and 13 HET). Unpaired t-test p= 3.9×10^−3^ (two-tailed, t= 3.15, df= 27). **(B)** Adult body weights (n=9 per genotype). Permuted Mann-Whitney U (MWU) p-values: wk4= 2×10^−4^, wk5 & 6< 1×10^−4^, wk7= 4×10^−4^, wk8= 3×10^−4^, wk9= 1.1×10^−3^, wk10< 1×10^−4^. Representative image of 8 wk adult WT and HET. **(C)** Vertical screen test. Permuted MWU p-values: P4= 0.31, P6= 0.05, P8= 0.07, P10= 0.07, P12= 0.96. **(D)** Surface righting test. Permuted MWU p-values: P4= 0.3, P6= 0.42, P8= 0.07, P10= 7×10^−4^, P12= 0.15. **(E)** 90° negative geotaxis test. Permuted MWU p-values: P4= 0.96, P6= 0.91, P8= 0.14, P10=0.16, P12= 4.2×10^−3^. **(F)** 180° negative geotaxis test. Permuted MWU p-values: P4= 0.39, P6=0.2, P8=0.16, P10=0.47, P12=0.08. (C-F, n=16 per genotype). PND= postnatal day. Error bars = SEM.

**Supplemental Figure 7.**
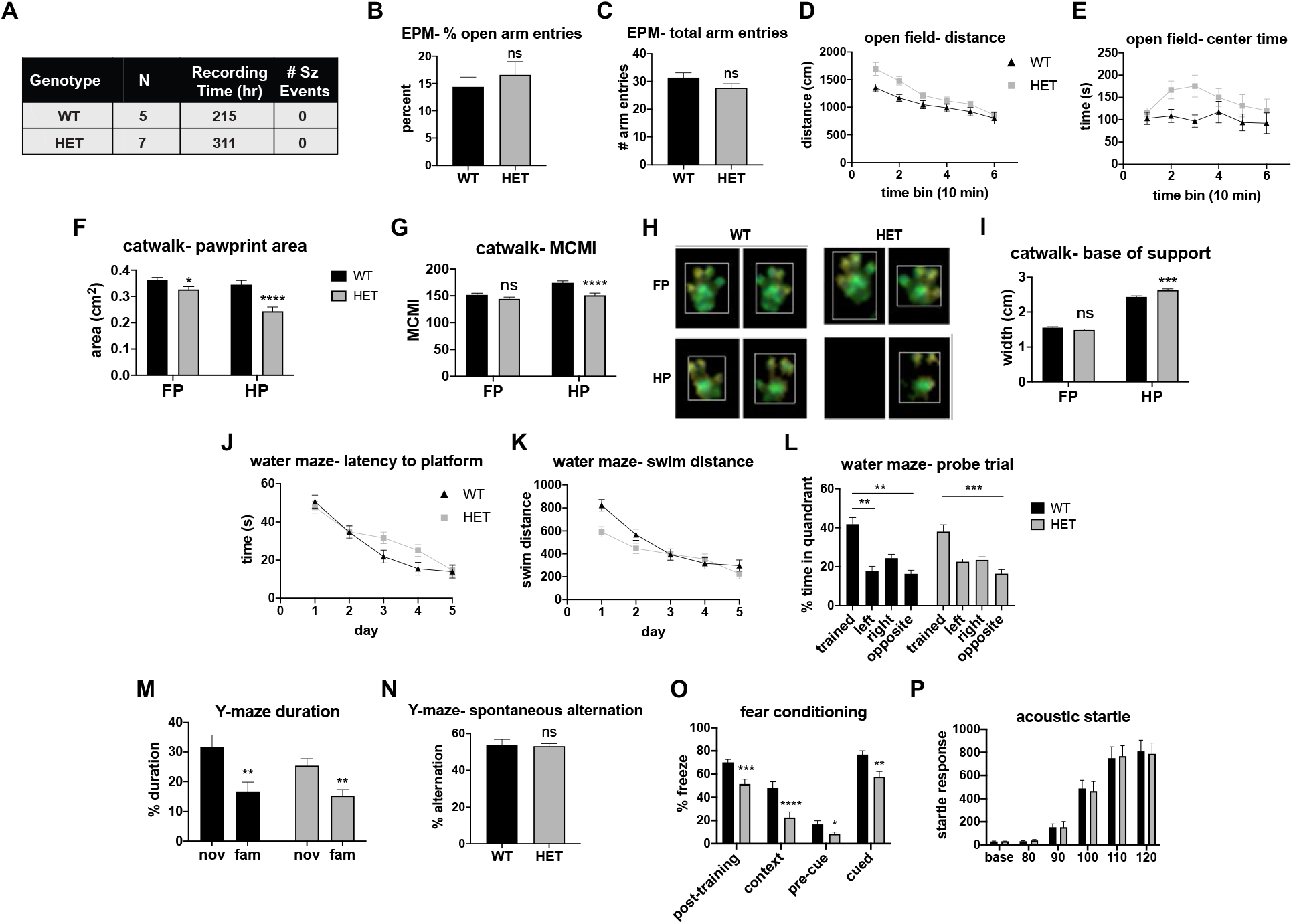
Adult behavior studies of *Hnrnpu*^+/113DEL^ mice. **(A)** Summary table of EEG data. Sz = seizure **(B)** Elevated plus maze (EPM) percent time spent in open arm (n= 14 WT, 22 HET). Welch’s t-test p= 0.47 (two-tailed, t= 0.73, df= 33.8). **(C)** EPM total arm entries (n= 14 WT, 22 HET). T-test p= 0.12 (two-tailed, t= 1.58, df= 34). **(D)** Open field ambulatory distance divided into six 10 min bins (n= 16 WT, 18 HET). Two-way repeated-measure (RM) ANOVA p= 0.026 (f= 5.4, df= 1). **(E)** Time spent in the center of open field, divided into 10 min bins (n= 16 WT, 18 HET). Two-way RM ANOVA p= 0.09 (f= 3.06, df= 1). **(F)** Catwalk front-paw (FP) and hind-paw (HP) print area (n= 17 WT, 19 HET). Unpaired, two-tailed t-test FP p= 0.024 (t= 2.4, df= 34), HP p< 1×10^−4^ (t= 4.4, df= 34). **(G)** Catwalk max contact max intensity (MCMI) of FP and HP (n= 17 WT, 19 HET). Mann-Whitney U (MWU) FP p= 0.08, HP p< 1×10^−4^. **(H)** Representative images of FP and HP prints of WT and HET mice. **(I)** Catwalk base of support for FP and HP (n= 17 WT, 19 HET). MWU FP p= 0.08, HP p= 1.5×10^−3^. **(J)** Morris Water Maze (MWM) latency to platform (acquisition period) measured over 5 consecutive days (n= 15 WT, 18 HET). Two-way RM ANOVA p= 0.22 (f= 1.56, df= 1). **(K)** MWM swim distance measured over 5 days (n= 15 WT, 18 HET). Two-way RM ANOVA p= 0.1 (f= 2.88, df= 1). **(L)** MWM probe trial measured as time spent in each quadrant of the pool (n= 15 WT, 18 HET). One-way RM ANOVA on ranks: WT T (trained quadrant) vs L (left quadrant) p< 1×10^−3^, T vs R (right quadrant) p= 0.2, T vs opp (opposite quadrant) p< 1×10^−3^. HET T vs R p= 2×10^−3^, T vs L p= 0.64, T vs opp p= 0.42. **(M)** Y-Maze percent duration spent in novel vs familiar arm (n= 16 WT, 18 HET). One-way RM ANOVA WT p= 5×10^−3^ (f= 10.5, df= 1), HET p= 3×10^−3^ (f= 11.5, df= 1). **(N)** Y-Maze percent spontaneous alternation (n= 16 WT, 18 HET). Welch’s t-test p= 0.86 (two-tailed, t= 0.18, df= 20.9). **(O)** Fear conditioning percent freezing (n= 16 WT, 18 HET). MWU post-train p= 7×10^−4^, context p< 1×10^−4^, pre-cue p= 0.02, cued p= 1.8×10^−3^. **(P)** Startle response (y-axis) across different decibels (x-axis) (n= 11 WT, 13 HET). T-test p= 0.96. Error bars= SEM.

**Supplemental Figure 8.**
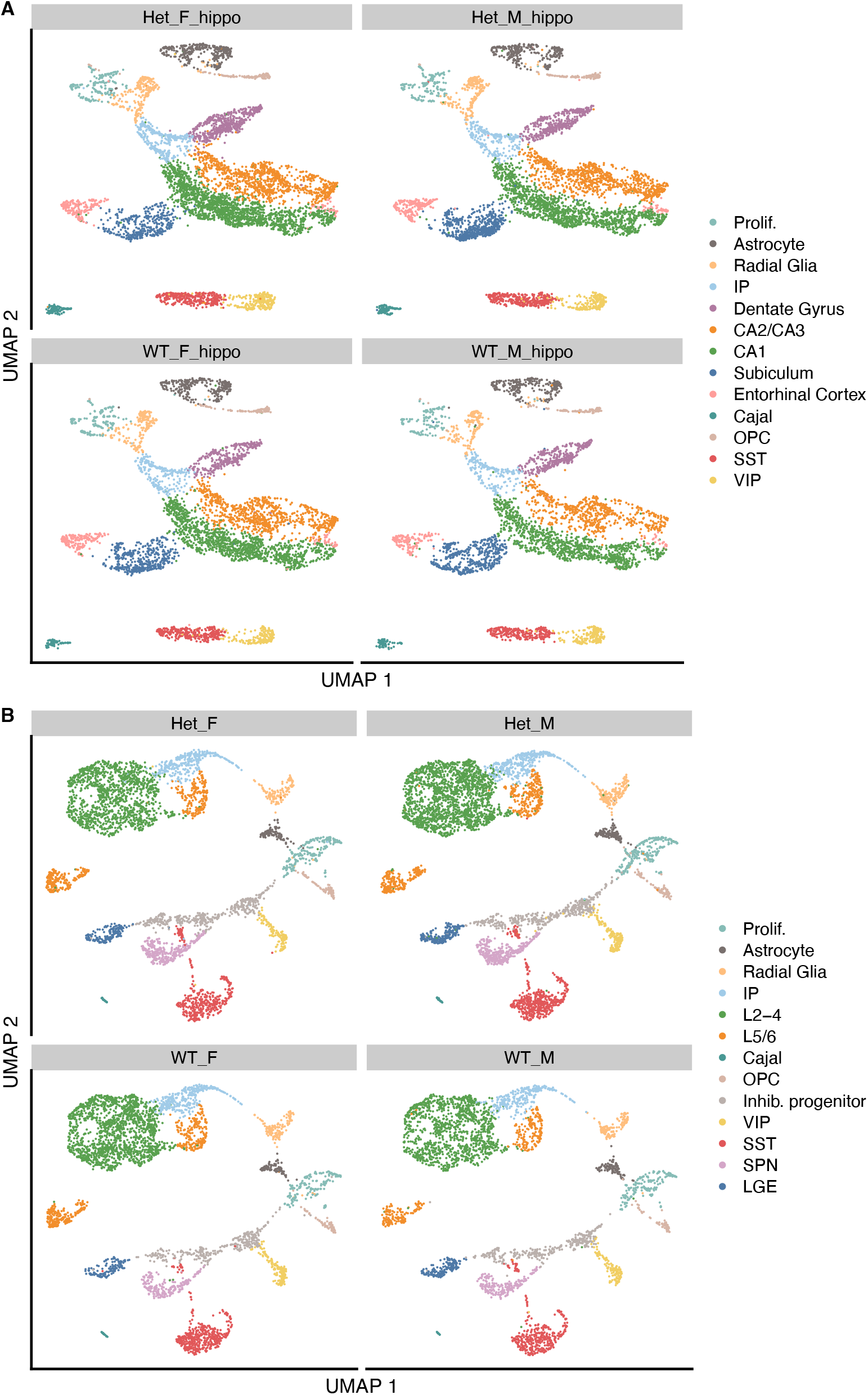
Single-cell RNA-seq dataset integration. **(A)** UMAP plots of cells from each hippocampal sample. **(B)** UMAP plots of cells from each cortical sample. Het_F: neocortical cells from a female; Het_M: neocortical cells from a male. Het_F= *Hnrnpu*^+/113DEL^ female; Het_M= *Hnrnpu*^+/113DEL^ male; WT_F= wildtype female; WT_M= wildtype male.

**Supplemental Figure 9.**
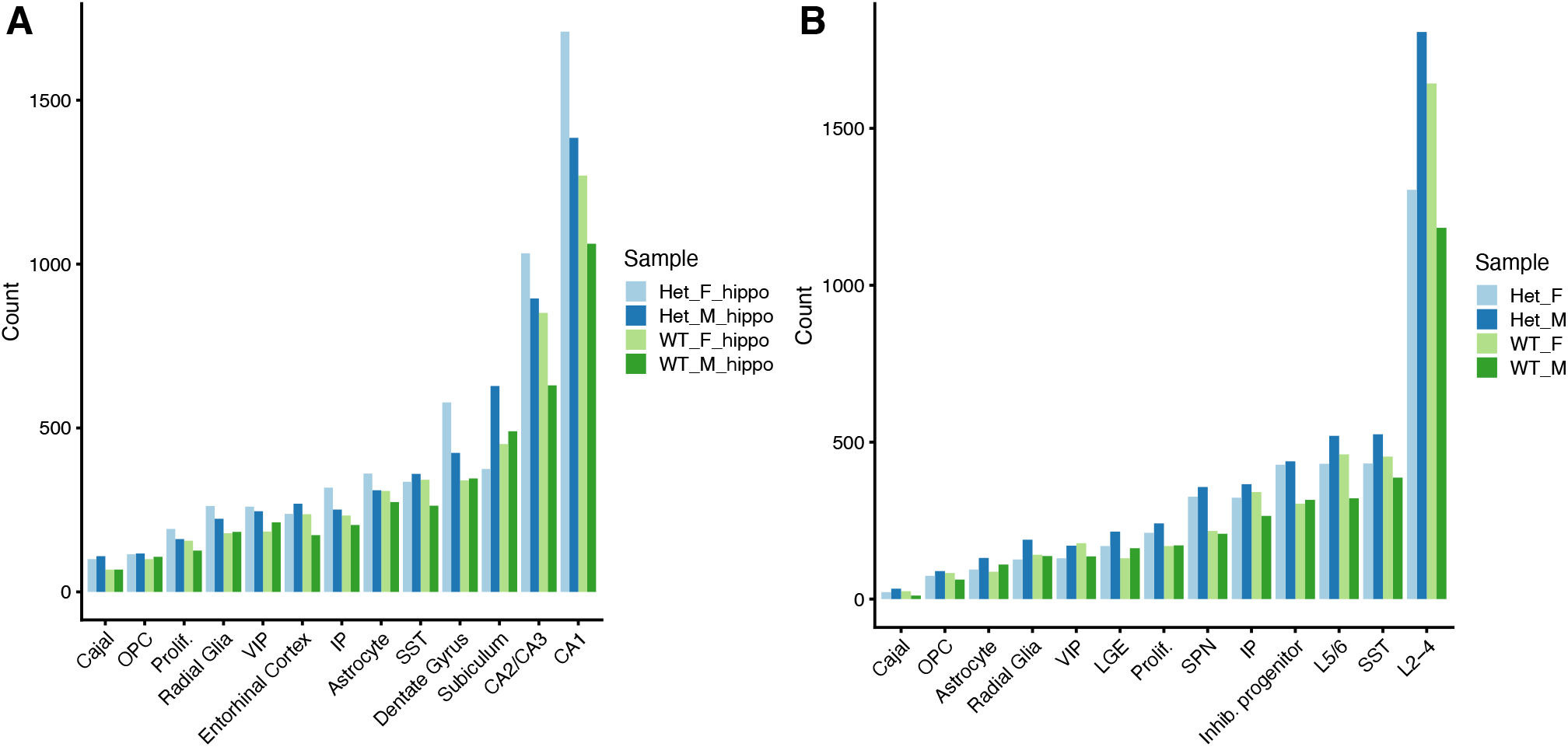
Number of cells per cluster. (**A**,**B)** Number of cells detected in each cell population, colored by sample. **(A)** represents hippocampal clusters and **(B)** represents neocortical clusters. Het_F= *Hnrnpu*^+/113DEL^ female; Het_M= *Hnrnpu*^+/113DEL^ male; WT_F= wildtype female; WT_M= wildtype male.

**Supplemental Figure 10.**
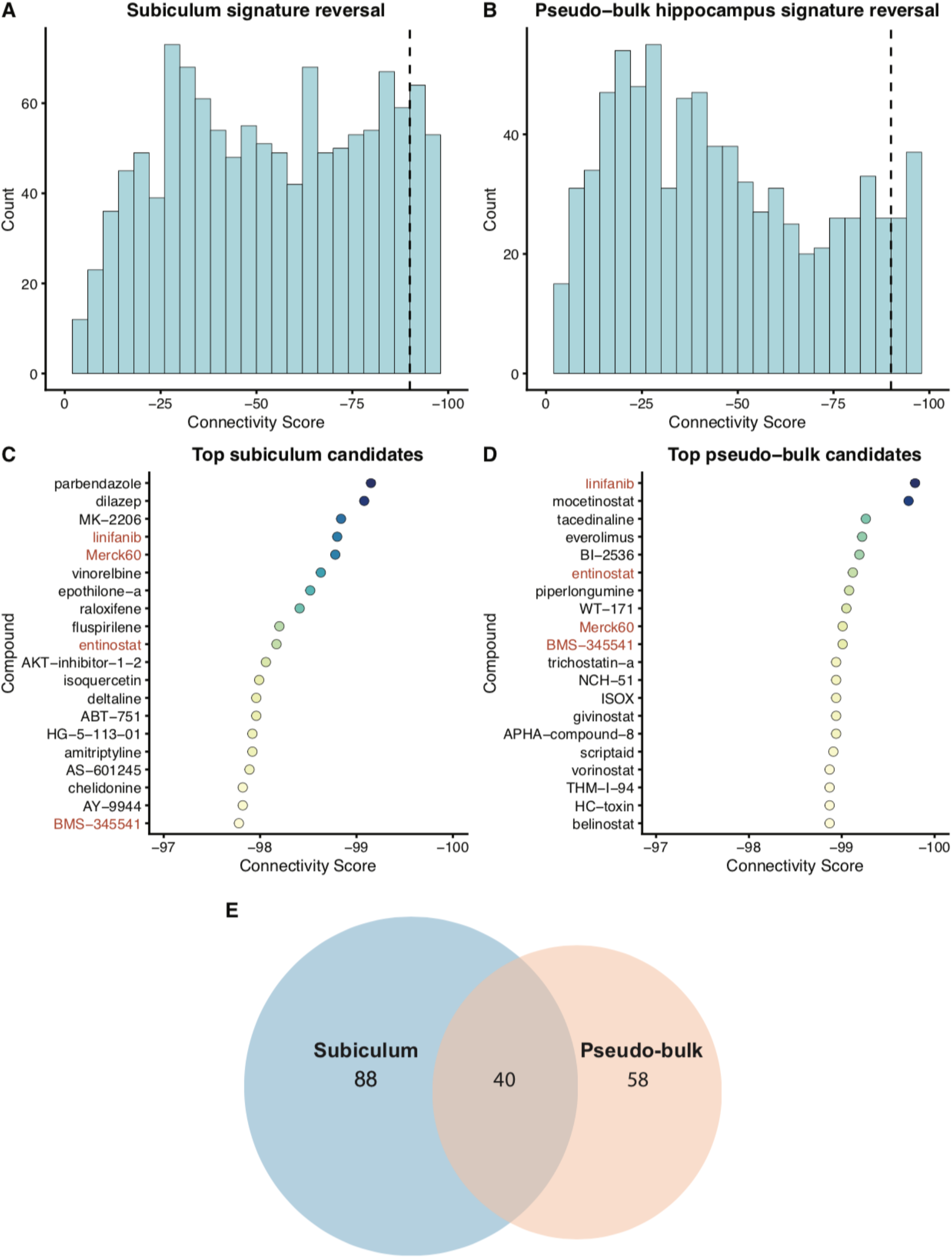
Transcriptomic signature reversal for hippocampal disease signatures. **(A, B)** Distribution of Connectivity Scores for the subiculum-derived and pseudo-bulk derived disease expression signatures. **(C, D)** Top 20 compounds predicted to reverse the subiculum and pseudo-bulk signatures. Compounds in red represent compounds prioritized for both signatures. **(E)** Venn Diagram depicting the overlap between all compounds achieving a Connectivity Score less than -90 for the subiculum and pseudo-bulk signatures.

## Additional supplemental files

File name: Supplemental Table 1

Description: List of published human *HNRNPU* pathogenic variants

File name: Supplemental Table 2

Description: Canonical cell-type-specific markers used to annotate cell clusters

File name: Supplemental Table 3

Description: Hippocampal differential gene expression results for all cell types

File name: Supplemental Table 4

Description: Neocortical differential gene expression results for all cell types

File name: Supplemental Table 5

Description: Gene ontology results for hippocampal and neocortical up and downregulated genes

File name: Supplemental Table 6

Description: Compounds with Connectivity Scores less than -90 for subiculum-derived and pseudobulk transcriptional signatures.

## Notes

https://www.ncbi.nlm.nih.gov/geo/query/acc.cgi?acc=GSE152715

### Summary of Updates

Included hIPSC-derived neuronal model qRT-PCR data and corresponding supplemental Fig 4; moved Fig 6 to the supplement; author affiliations updated; minor text edits

