## Supplemental Table 1 for "Neurodevelopmental deficits and cell-type-specific transcriptomic perturbations in a mouse model of *HNRNPU* haploinsufficiency"

| cDNA change (NM_031844.2) | Amino Acid change | Mutation Type | Inheritance | Reference |
|---|---|---|---|---|
| c.16delins | p.Val6Ilefs*4 | PPT | Paternal mosaicism | Depienne et al., 2017 |
| c.23del | p.Val8Glufs*4 | PPT | De novo | Yates et al., 2017; |
| c.418G>A | p.Glu140Lys | Missense | De novo | Yates et al., 2017 |
| c.511C>T | p.Gln171* | PPT | De novo | Hamdan et al., 2014 |
| c.523C>T | p.Gln175* | PPT | De novo | Bramswig et al., 2017 |
| c.651_660del | p.Gly218Alafs*118 | PPT | De novo | Leduc et al., 2017 |
| c.692-1G>A | p.? | Splice site | De novo | Depienne et al., 2017; |
| c.817C>T | p.Gln273* | PPT | De novo | Bramswig et al., 2017 |
| c.960G>A | p.Trp320* | PPT | De novo | Yates et al., 2017; |
| c.970A>G | p.Arg324Gly | Missense | De novo | Bramswig et al., 2017 |
| c.1089G>A | p.Trp363* | PPT | De novo | Leduc et al., 2017 |
| c.1117+1G>A | p.? | Splice site | De novo | Yates et al., 2017; |
| c.1132T/C | p.Ser378Pro | Missense | De novo | Bramswig et al., 2017 |
| c.1424_1425insTC | p.Ile476Profs*7 | PPT | De novo | Yates et al., 2017; |
| c.1615-1G>A | c.1615-1G>A | Splice site | De novo | Need et al., 2012; Zhu et al. 2015 |
| c.1626_1627insA | p.Lys543* | PPT | De novo | Yates et al., 2017; |
| c.1664del | p.Leu555Argfs*51 | PPT | De novo | Yates et al., 2017; |
| c.1681C>T | p.Gln561* | PPT | De novo, mosaic | Depienne et al., 2017 |
| c.1681del | p.Gln561Serfs*45 | PPT | De novo | Depienne et al., 2017 |
| c.1714C>T | p.Arg572* | PPT | De novo | Leduc et al., 2017 |
| c.1744-4_1749del | p.? | Splice site | De novo | Epi4K Consortium et al. 2013 |
| c.1868dup | p.Glu624Argfs*24 | PPT | De novo | Depienne et al., 2017; de Kovel et al., 2016 |
| c.2270_2271del | p.Pro757Argfs*7 | PPT | De novo | Leduc et al., 2017 |
| c.2299_2302del | p.Asn767Glufs*66 | PPT | De novo; mosaic | Depienne et al., 2017 |
| c.2425-3C>A | p.? | Splice site | De novo | Depienne et al., 2017 |
| c.2471_2472delinsGA | p.Tyr824* | PPT | Unknown; mother negative | Carvill et al., 2013 |
